# Electro-Osmotic Flow Generation via a Sticky Ion Action

**DOI:** 10.1101/2023.12.14.571673

**Authors:** Behzad Mehrafrooz, Luning Yu, Zuzanna Siwy, Meni Wanunu, Aleksei Aksimentiev

## Abstract

Selective transport of ions through nanometer-sized pores is fundamental to cell biology and central to many technological processes such as water desalination and electrical energy storage. Conventional methods for generating ion selectivity include placement of fixed electrical charges at the inner surface of a nanopore through either point mutations in a protein pore or chemical treatment of a solid-state nanopore surface, with each nanopore type requiring a custom approach. Here, we describe a general method for transforming a nanoscale pore into a highly selective, anion-conducting channel capable of generating a giant electro-osmotic effect. Our molecular dynamics simulations and reverse potential measurements show that exposure of a biological nanopore to high concentrations of guanidinium chloride renders the nanopore surface positively charged due to transient binding of guanidinium cations to the protein surface. A comparison of four biological nanopores reveals the relationship between ion selectivity, nanopore shape, composition of the nanopore surface, and electro-osmotic flow. Remarkably, guanidinium ions are also found to produce anion selectivity and a giant electro-osmotic flow in solid-state nanopores via the same mechanism. Our sticky-ion approach to generate electro-osmotic flow can have numerous applications in controlling molecular transport at the nanoscale and for detection, identification, and sequencing of individual proteins.

## Introduction

Ion transport across biological membranes is integral to numerous physiological functions, including osmotic balance maintenance, electrical signaling, cell volume regulation, cell-to-cell communication, and muscle contraction.^1^ Such ion transport is enabled by a diverse family of membrane proteins that act as pumps,^2,3^ transporters^4^ and ion channels,^5,6^ providing a conduit across otherwise impermeable membranes. The function of ion channels, however, is not limited to serving as a passive ion passageway—they also act as ion traffic controllers, regulating the direction of the ion transport and selectively conducting specific ion species. ^7^

Crystallographic studies have unraveled the detailed molecular architecture of multiple ion channels, revealing the intricate mechanisms underlying their ion transport and selectivity function. Armed with the wealth of such structural information, on-going efforts are directed to alter ion transport functionality of biological channels through protein engineering^8^ or design completely new protein channel architectures.^9,10^ Common approaches to tune ion selectivity include altering the physical size of the channel’s constriction *via* a steric exclusion mechanism,^11^ or placing fixed charges at the nanopore surface to create a selective electrostatic barrier. ^12^

While many biological channels exhibit superb ion selectivity, their practical applications are tempered by the elaborate process of their extraction from a biological sample, the nearly fixed shape and size of their transmembrane pores, as well as their mechanical fragility and sensitivity of their performance to environmental factors such as pH and temperature. ^13^ Consequently, a large body of work has been directed toward reproducing the biological performance of ion channels in more robust synthetic material systems.^14^ Ion selectivity was purposefully engineered in solid-state nanopores by customizing the nanopore geometry, ^15–17^ by chemically modifying the nanopore surface,^18–22^ and by introducing asymmetric surface charge distributions.^23–28^, or even mechanical strain. ^29^ Although these synthetic solid-state nanopores do offer superior mechanical and chemical stability and are amenable to custom shape and charge modifications,^30,31^ they usually have subpar spatial and temporal resolutions in biosensing applications,^32–36^ suffer from pore clogging issues, ^37^ and their fabrication remains relatively expensive.

One particular application of transmembrane pores is in nanopore sensing and biopolymer sequencing, where the chemical identity of a biological molecule is determined by recording the ionic current blockade signature produced by during molecular passage.^38–40^ Transport of charged molecular species can be achieved by the application of a transmembrane voltage, however, the majority of biological molecules—proteins in particular—have a heterogeneous distribution of their electrical charge. Although mechanical pulling a polypeptide chain through a nanopore can be realized using an AFM tip^41–44^ or a motor protein,^41^ pushing biomolecules through a nanopore by means of a water flow has emerged as the most versatile and convenient method for achieving the transmembrane transport of analytes with hetergeneous charge distributions.^45–47^ The most common and convenient method of producing such a flow is by utilizing the electro-osmotic effect. ^48,49^ When a voltage is applied across an ion-selective nanopore, the selectivity of the ionic current results in a unidirectional fluid motion known as the electro-osmotic flow (EOF). Generating and harnessing such EOF is in particular important for controlling protein transport in single-molecule protein sequencing, the current frontier of nanopore sensing applications. ^50^

Here we describe a general method of generating EOF in a nanopore by transforming the nanopore into an anion-selective channel by the addition of guanidinium chloride (GdmCl). Our molecular dynamics (MD) simulations and reverse potential measurements show that the presence of guanidinium produces strong chloride transport which is caused by Gdm^+^ ions binding to the pore walls. We then show how such ion selective current translates into EOF generation and demonstrate the generality of the effect by examining four biological and one solid-state nanopore. Our results set the stage for rational engineering of nanopores for electro-osmosis-assisted transport of analytes, irrespective of their charge distributions.

## Results

While measurements of ionic current are readily accessible to experiments, experimental characterization of EOF is difficult and indirect^51,52^ as it relies on the motion of a tracer particle.^45,53,54^ In contrast, all-atom MD simulations can directly characterize the EOF by tracking the motion of each water molecule in response to the application of the external electric field.^55,56^ In what follows, we first examine the performance of the all-atom MD method in predicting the ionic selectivity of a biological nanopore and then apply this computational method to characterize EOF and its origin across multiple pores.

### Ion selectivity of *α*-hemolysin from simulation and experiment

As a test system for our computational method, we chose *α*-hemolysin— a bacterial toxin widely used in nanopore translocation experiments. ^57^ Figure 1a shows an all-atom model of the experimental system containing one *α*-hemolysin nanopore, a patch of a lipid membrane and 2.5 M KCl electrolyte solution. After a brief equilibration using the all-atom MD method, the ionic current through *α*-hemolysin was simulated under a 200 mV transmembrane bias of either polarity using a previously established protocol. ^55^ Figure 1b plots the total charge carried by each ion species through the transmembrane pore of the protein over the course of the MD simulation under +200 mV. The slope of the charge transfer curve, *i*.*e*., the average current, is about 2.5 times greater for chloride than for potassium, indicating that the current is anion selective. Under a reversed bias polarity (−200 mV), the ionic current remains anion-selective, although the selectivity is reduced (Figure 1c and Figure S1a).

**Figure 1:**
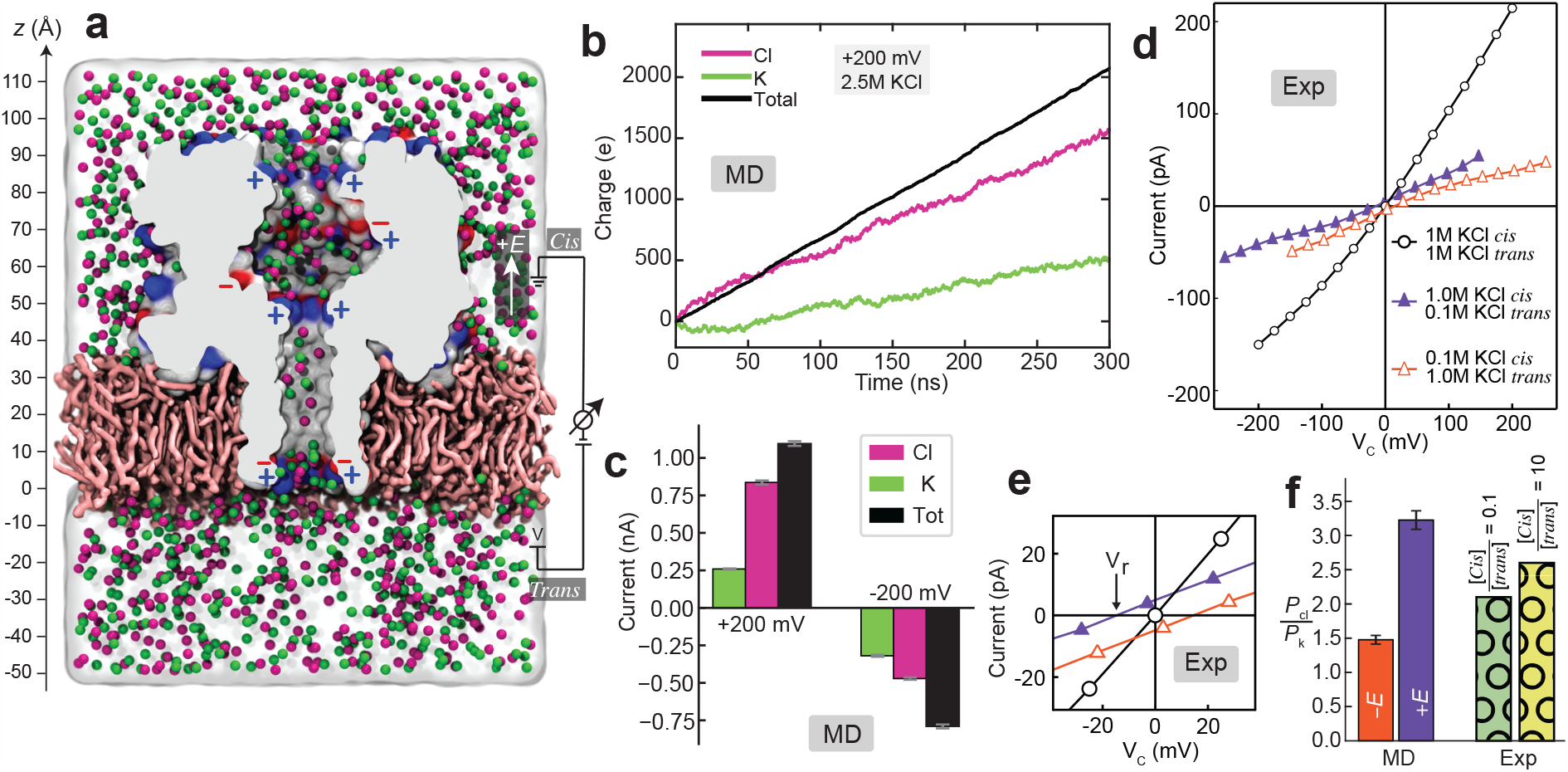
Ion selectivity of *α*-hemolysin in KCl electrolyte. **a**, All-atom model of *α*-hemolysin (cutaway molecular surface) embedded in a POPC membrane (pink) and submerged in 2.5 M KCl electrolyte (white semitransparent surface; K^+^ and Cl^−^ ions shown as green and magenta spheres, respectively). The surface of *α*-hemolysin is color-coded according to the charge of the surface residues, blue for positive, red for negative and white for neutral residues. **b**, Total charge carried by ion species through the transmembrane pore of *α*-hemolysin over the course of an MD simulation under +200 mV bias. The slope of each line corresponds to the average current. **c**, Average simulated current carried by the ionic species at +/−200 mV. Negative values indicate transport in the direction opposite that of the *z*-axis (defined in panel a). Hereafter, the average current is computed as the slope of the respective charge-time curve and the error bars show the counting error of mean computed from the total number of transported ions. **d**, Experimentally measured current-voltage (*I* -*V*) curves of *α*-hemolysin obtained at three gradients of ion concentration. The *cis* and *trans* compartments are defined in panel a. The voltage, *V*_c_, has been corrected to account for the Nernst potential at the electrodes. ^61^ The uncorrected *I*-*V* curves are shown in Figure S2. **e**, Zoomed-in view of panel d. A reversal potential, *V*_r_, is determined as the *x*-intercept of the *I*-*V* curves. **f**, Simulated (MD) and experimentally determined (Exp) ratios of ion species permeability through *α*-hemolysin at two polarities of the transmembrane bias (MD) and at two polarities of the concentration gradient (Exp). All experimental measurements were performed at 25^*°*^ C and 10 mM Tris buffering the pH at 7.5.

Complimenting MD simulations, we measured ion selectivity of *α*-hemolysin under various experimental conditions. Under typical nanopore measurement conditions, *e*.*g*. when 1 M KCl electrolyte solution is placed on both sides of the membrane, the *α*-hemolysin channel exhibits a voltage-dependent ion conductance (Figure 1d), which originates from the asymmetric shape of the channel and the non-uniform charge distribution along its transmembrane pore.^58,59^ By comparing the simulated ionic currents with their respective experimental values (Table S1), we find the MD method to systematically overestimate the simulated ionic current magnitudes, which can be largely attributed to the differences in the simulated and experimental bulk electrolyte conductivities (Table S2). Nevertheless, the ratio of the ionic currents simulated at transmembrane biases of the same magnitude but opposite polarities, *i*.*e*., the ionic current rectification, matches quantitatively the experimentally measured values, in accordance with the previous studies.^55,60^

To experimentally characterize the ion selectivity of *α*-hemolysin, we repeated current *versus* voltage measurements under asymmetric electrolyte conditions, where the electrolyte concentration on the one side of the membrane was ten times that on the other side (Figure 1d). From the recorded *I*-*V* curves, we determined a so-called reversal potential, *V*_r_, which is the voltage at which the transmembrane current is zero. For the 1 M [KCl]_*cis*_ */* 0.1 M [KCl]_*trans*_ electrolyte condition, the reserve potential is measured to be −14.6 mV (Figure 1E), which corresponds to Cl^−^ permeability being about twice that of K^+^, *i*.*e*., *P*_Cl_*/P*_K_ = 2.1, please see SI Material and Methods for the detailed description of the calculations. Interestingly, for the reverse asymmetric condition of 0.1 M [KCl]_*cis*_ / 1 M [KCl]_*trans*_, we find that *P*_Cl_*/P*_K_ = 2.6 (Figure 1f). The 25% higher selectivity observed when the vestibule (*cis*) side is exposed to a lower salt concentration is possibly caused by the weaker electrostatic screening of the vestibule charge (Figure 1a) by the counterions. This observation suggests that the positively charged vestibule of *α*-hemolysin is largely responsible for the channel’s anion selectivity in KCl electrolyte. ^62^

The measured selectivity is in accordance with the results of prior studies^62,63^ which found the ratio of Cl^−^ to K^+^ currents to be about 2:1, despite both ions having very similar electrophoretic mobilities. The results of the measurements are also in good agreement with the ion selectivity determined directly from the all-atom MD simulations: *P*_Cl_*/P*_K_= 1.5 *±* 0.2 and 3.2 *±* 0.4 for the *cis*-to-*trans* and the *trans*-to-*cis* directions of the electric field (Figure 1f). Note that a direct quantitative comparison is not possible as the MD simulations assess ion selectivity under the equal concentration electrolyte conditions, the conditions typically realized in nanopore translocation measurements.

### Guanidinium transforms *α*-hemolysin into a highly selective anion channel

To investigate the effect of GdmCl on ion selectivity of *α*-hemolysin, we modified our simulation system to contain 1.5 M of GdmCl electrolyte (Figure 2a). Figure 2b illustrates the transport of charged species through the nanopore when the system was simulated under a +200 mV bias. The ionic current through the nanopore (the slope of the charge *versus* time plot) is found to be carried mostly by the chloride ions whereas the current carried by guanidinium ions was close to zero. Furthermore, when the simulations were repeated using a mixture of GmdCl and KCl (1.5 M GdmCl / 1.0 M KCl), the ionic current through the nanopore not only remained anion selective, but also the current carried by potassium ions was close to zero (Figure 2c and Figure S1b). Strong anion selectivity was also observed for the reversed polarity of the transmembrane bias (Figure 2c and Figure S1c-d).

**Figure 2:**
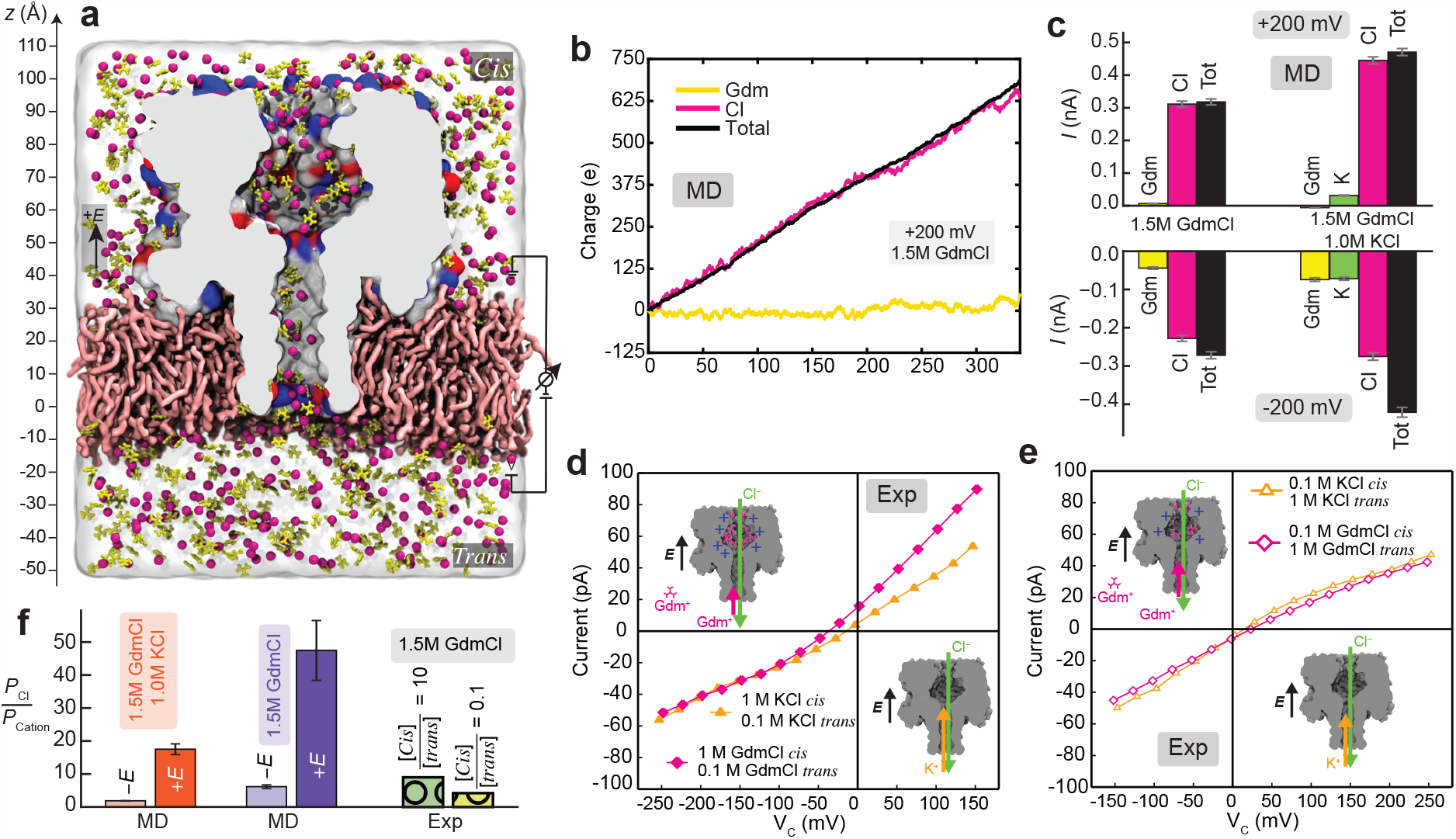
Ion selectivity of *α*-hemolysin in the presence of guanidinium. **a**, All-atom model of *α*-hemolysin submerged in 1.5 M guanidinium chloride electrolyte. Guanidinium ions are shown in yellow, all other elements of the system are shown as in Figure 1a. **b**, Total charge carried by ion species through the transmembrane pore of *α*-hemolysin over the course of an MD simulation under +200 mV. **c**, Simulated average current carried by the ionic species through *α*-hemolysin at +/−200 mV for a pure 1.5 M GmdCl electrolyte and a mixture of 1.5 M GmdCl and 1 M KCl. **d**,**e** Experimentally measured current-voltage dependence of *α*-hemolysin for two concentration gradients of GmdCl. Data for similar gradients of KCl are shown for comparison. The reversal potential, *V*_*r*_, is measured as the *x* -intercept of the *I*-*V* curves. **f**, Experimental and simulated ratios of the ionic species permeability at two polarities of the transmembrane bias (MD) and two concentration gradients (Exp). All experimental measurements were performed at 25^*°*^ C and 10 mM Tris buffering the pH at 7.5.

Complementing our MD simulations, the ion selectivity of *α*-hemolysin in the presence of guanidinium ions was characterized experimentally. Figures 2d and 2e show the *I*-*V* curves measured for the 1 M [GdmCl]_*cis*_ / 0.1 M [GdmCl]_*trans*_ and 0.1 M [GdmCl]_*cis*_ / 1 M [GdmCl]_*trans*_ electrolyte conditions, respectively. *I*-*V* curves for equivalent asymmetric KCl conditions are shown for comparison. The presence of guanidinium makes the non-linear character of the curves more pronounced. The reverse potential values extracted from the curves are −34.5 mV and 25.0 mV, which translates into a *P*_Cl_*/P*_Gmd_ ratios of 9.0 and 4.2 for the two polarities of the concentration gradient (Figure 2f). That is, at low concentrations of GdmCl in the *cis* (vestibule) chamber, *α*-hemolysin shows weaker Cl^−^ selectivity as compared to the case of a higher GdmCl concentration. Thus, the asymmetry of the channel selectivity in the two directions of the GdmCl gradient is opposite to that determined for KCl (compare Figure 1f and Figure 2f). This observation is surprising and suggests that the ion selectivity of *α*-hemolysin in the presence of GdmCl cannot be explained by the electrostatic screening argument.

The high anion selectivity of *α*-hemolysin experimentally measured for 1.5 M GmdCl electrolyte corroborates the high anion selectivity observed directly from MD simulations (Figure 2f). Note the high statistical error in calculating the Gdm^+^ current from the MD trajectory at +200 mV, as only a few Gdm^+^ ions were observed to pass through the nanopore over the simulation time scale. Finally, we note that, in dilute solutions (10 mM), the measured bulk conductivities of KCl (1.47 mS/cm) and GdmCl (1.21 mS/cm) differ by only∼20%, which suggests that electric mobilities of Gdm^+^ and K^+^ are not very different. This conclusion is in line with the previously determined diffusion coefficients of the two ion species: 2.15 *×* 10^−9^ m^2^/s for Gdm^+^ and 1.86 *×* 10^−9^ m^2^/s for K^+^.^64,65^ The relatively small difference in the mobility of the two cations cannot account for the large anion selectivity obtained when switching from KCl to GdmCl electrolyte.

### Guanidinium-induced anion selectivity is universal across biological nanopores

To determine the generality of the guanidinium-induced anion selectivity, we built and simulated three other biological nanopore systems. The first system included the M1-NNN mutant of MspA^66^ (Figure 3a), a nanopore widely used in development of DNA and protein sequencing methods. Three variants of the system were built differing by the concentration and composition of the electrolyte: pure 2.5 M KCl, pure 1.5 M GdmCl, and a mixture of 1.5 M GdmCl / 1.0 M KCl. Each system was simulated under a 200 mV bias of either polarity. Among the three systems, the highest absolute conductance was recorded for the pure KCl electrolyte system, which also exhibited a very mild ion selectivity: 51.8% of the current was carried by Cl^−^ ions (Figure 3b and Figure S3). Replacing K^+^ with Gdm^+^ reduced the total ionic current, while simultaneously making the current anion selective, *P*_Cl_*/P*_Gdm_ = 2.3 *±* 0.08 at 1.5 M GdmCl. Same anion-selectivity, *P*_Cl_*/P*_Gdm+K_ = 2.2 *±* 0.1 was observed for the mixed composition of the electrolyte.

**Figure 3:**
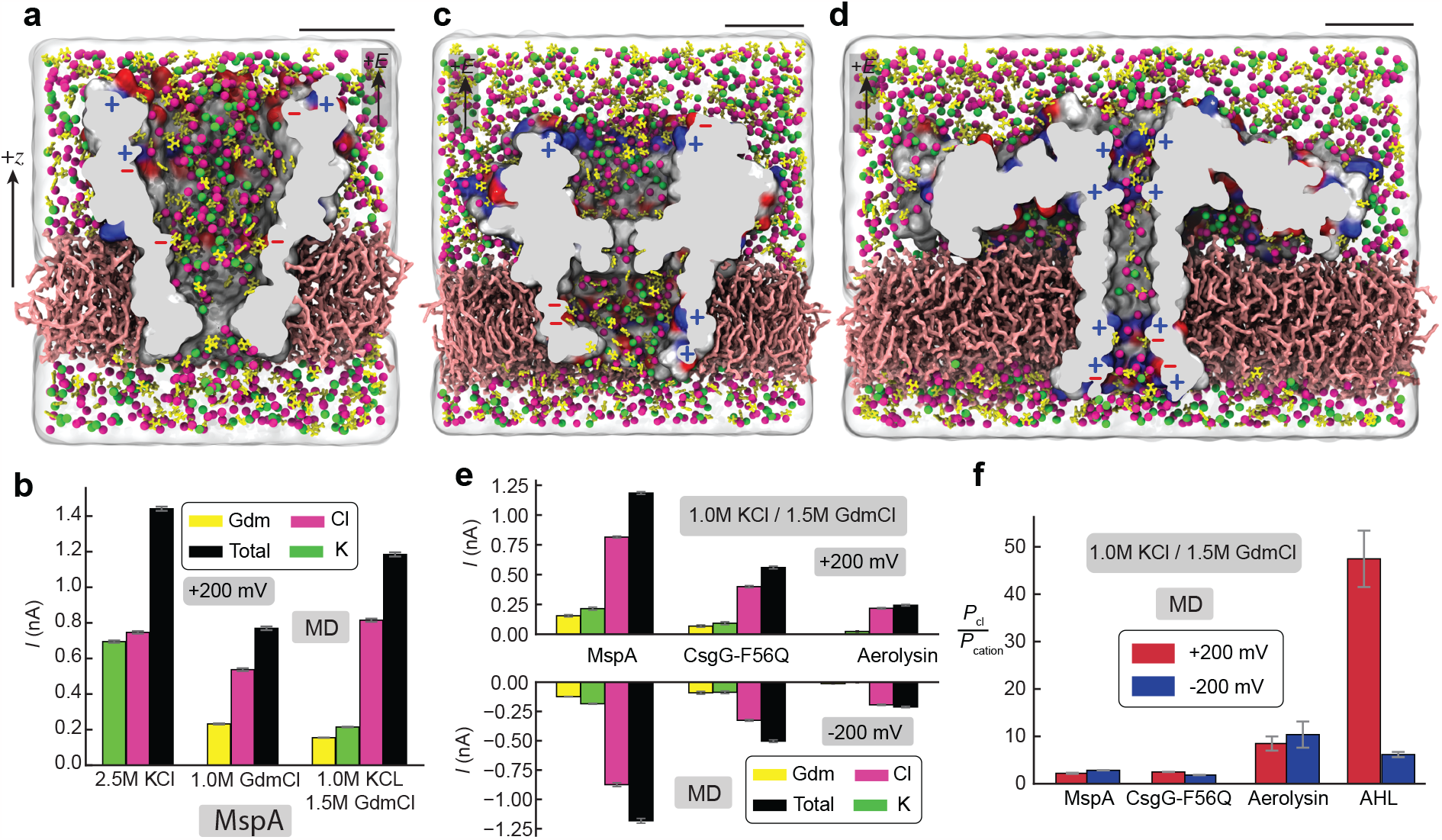
MD simulation of guanidium-induced anion selectivity in different biological nanopores. **a**, All-atom model of M1-NNN MspA embedded in a lipid bilayer membrane (pink) and surrounded by a 1.5 M GdmCl / 1.0 M KCl electrolyte mixture shown using a cut-away representation. Positively and negatively charged residues at the nanopore surface are shown in red and blue, respectively. Semitransparent surface illustrates the volume occupied by the electrolyte. Gdm^+^, K^+^ and Cl^−^ ions are colored in yellow, green, and magenta, respectively. The scale bar corresponds to 3 nm. **b**, Average simulated ionic current through MspA under +200 mV for the three electrolyte conditions. **c**,**d**, All-atom model of F56Q mutant of CsgG (panel c) and of wild type aerolysin (panel d). **e**, Ionic current carried by different ion species in MD simulations of the three nanopores under *±* 200 mV. **f**, Simulated ion selectivity of the four biological nanopores characterized as the ratio of anion current (Cl^−^) over the cation current (Gdm^+^ + K^+^).

The other two biological nanopore systems contained either an F56Q mutant of CsgG (Figure 3c)—the pores used in a commercial nanopore sequencing platform, ^67^ or a wild type aerolysin channel (Figure 3d)—a pore widely used for peptide sensing experiments. ^68,69^ Each system was built to contain a 1.5 M GdmCl / 1.0 M KCl electrolyte mixture. Under the same transmembrane bias, the total current flowing through the CsgG nanopore was smaller than through MspA and even smaller currents were recorded for aerolysin with the majority of the current being carried by anions (Figure 3e, Figure S4a-c, and Figure S5a-c). Under the same electrolyte conditions, the ion selectivity ratios extracted from the simulations of MspA and CsgG were comparable (∼2), but much lower than the selectivity ratios observed for aerolysin and *α*-hemolysin (Figure 3f). Thus, the anion selectivity appears to be correlated with the nanopore geometry, with the sequencing pores having a sharp constriction (MspA and CsgG) exhibiting less selectivity than the nanopores used for whole molecule sensing (aerolysin and *α*-hemolysin), which feature a long and narrow beta-barrel.

### Binding of Gdm^+^ ions to nanopore surface generates anion selectivity

In search of a mechanism responsible for the anion selectivity produced by guanidinium, we first examined the distribution of the electrostatic potential within *α*-hemolysin at the three electrolyte conditions (Figure 4a). The presence of guanidinium was found to produce rather minor differences in the average electrostatic potential along the nanopore axis, making local features of the potential profile less sharp. Similarly close-to-negligible effects on the average electrostatic potential were previously reported from a comparative study of five monovalent alkali electrolytes. ^60^

**Figure 4:**
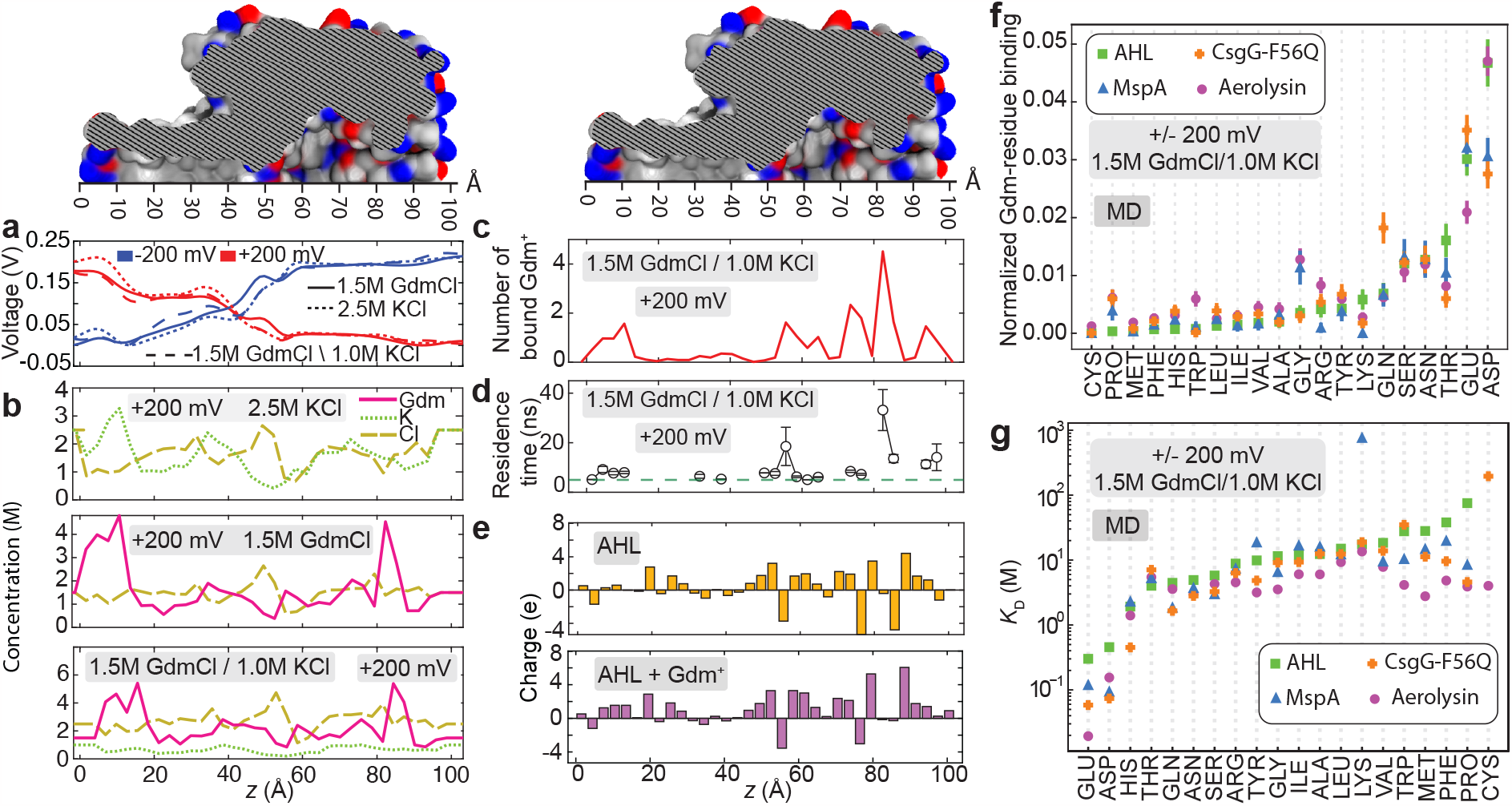
Guanidinium binds to nanopore surface and renders it positively charged. **a**, Average electrostatic potential along the axis of the *α*-hemolysin nanopore (the *z* -axis) for the three electrolyte conditions and at *±*200 mV transmembrane bias. **b**, Local concentration of ions along the transmembrane nanopore of *α*-hemolysin observed in all-atom MD simulations under +200 mV and 2.5 M KCl (top), 1.5 M GdmCl (middle) and 1.5 M GdmCl / 1.0 M KCl (bottom) electrolytes. The concentrations were averaged over the respective MD trajectories in 3 Å bin along the nanopore axis. **c**, Number of Gdm^+^ ions bound to the inner surface of the *α*-hemolysin nanopore, averaged over the simulation time. A Gdm^+^ ion was considered bound if any of its non-hydrogen atoms was located within 3 Å of any non-hydrogen atom of the nanopore. **d**, Average residence time of the Gdm^+^ ions bound to the *α*-hemolysin surface as a function of their location within the nanopore. Contacts between Gdm^+^ and *α*-hemolysin lasting less than 5 ns (dashed green line) were discarded from the residence time analysis. **e**, Local charge of the inner surface of the *α*-hemolysin nanopore (top) and the total charge of that surface when taking into account bound Gdm^+^ ions (bottom). **f**, Number of solvent-exposed residues of the specified type that bind Gdm^+^ normalized to the total number of such residues for the four biological nanopores. Error bars show the standard error computed using 30 ns fragments of the simulation trajectories. **g**, Dissociation constant (*K*_D_) computed for each type of amino acids in four different nanopores. Note the logarithmic scale of the *K*_D_ axis. In panels f and g, amino acid types are arranged in ascending order according to *α*-hemolysin data.

The presence of guanidinium was found to affect partitioning of ions into the nanopore. In the case of 2.5 M KCl, the local concentrations of K^+^ and Cl^−^ within the nanopore volume (Figure 4b) are similar, averaging to 1.4 M and 1.5 M, respectively. Similar average concentrations are seen in the case of 1.5 M GdmCl electrolyte (1.6 M for Gdm^+^ and 1.5 M for Cl^−^), despite the 1.6-times difference in the bulk electrolyte concentrations. In the mixed electrolyte system (1.5 M GdmCl / 1.0 M KCl), however, the concentration of K^+^ ions is three times lower than that of Gdm^+^, indicating that Gdm^+^ ions expel K^+^ ions out from the nanopore. In all systems, the concentrations of cations and anions within each section of the nanopore are largely similar (Figure 4b), thus satisfying the local electroneutrality condition. The ion concentration profiles were found to be insensitive to the polarity of the transmembrane bias (Figure S6).

According to Figure 4c, Gdm^+^ ions accumulate in the following three regions: near the *trans* exit of the nanopore stem, above the stem-vestibule junction and near the *cis* exit from the *α*-hemolysin vestibule (see Figure S7a-c for other electrolyte compositions and transmembrane biases). In the same three regions, individual Gdm^+^ ions remain bound to the nanopore surface for considerable (*>* 5 ns) intervals of time, Figure 4d and Figure S7d-f. Such binding of Gdm^+^ adds about 25 proton charges, on average, to the nanopore surface (Figure 4e). That surface charge is compensated by much more mobile chloride ions that carry the majority of the ionic current through the nanopore. The local binding of Gdm^+^ to the inner nanopore surface mildly correlates with the local charge of the nanopore, Figure 4e. Despite the applied bias, the number of Gdm^+^ ions bound to the nanopore surface remains largely constant (Figure S8).

The above ion binding mechanism can explain the unexpected reversal of the ion selectivity magnitude measured experimentally using concentration gradients of KCl and GdmCl. In the case of KCl, a higher anion selectivity was measured when a lower concentration of KCl was in the *α*-hemolysin vestibule (Figure 1f), which is explained by a weaker screening of the fixed charge of the vestibule residues. The opposite was found for GdmCl: a higher anion selectivity was measured at a higher concentration of Gdm^+^ in the vestibule (Figure 2f), which is consistent with more Gdm^+^ ions binding to the surface of the vestibule as more such ions are present in the solution.

To quantitatively characterize the local affinity of guanidinium to the nanopore surface, we computed the average number of contacts made by Gdm^+^ ions with the residues of the protein surface for each type of the twenty amino acid types, normalizing each number by the abundance of such surface residues. The resulting ranking of local Gdm^+^ affinity was found to be insensitive to the polarity of the transmembrane bias (Figure S9). Using the data at both bias polarities, we ranked Gdm^+^ affinity to the twenty amino acids in the four biological nanopores (Figure 4f). Among the twenty amino acids, the negatively charged residues (aspartic acid and glutamic acid) were found to be the most favorable sites for guanidinium binding, followed by residues that could exhibit hydrophobic interactions. Interestingly, the positively charged residues (lysine and arginine) were not at the bottom of the ranking. Overall, the ranking was consistent across the four biological nanopores. We attribute the deviations from the average ranking to a rather large steric footprint of Gdm^+^ (ten atoms, ∼ 4 *×* 4 Å^2^), which could simultaneously contact more than one surface residue.

Using the framework of a ligand-enzyme kinetic analysis, we computed the dissociation coefficients, *K*_D_, for guanidinium and the biological nanopores, see SI for details. Assuming all surface residues of the protein to have the same binding affinity, we find the average *K*_D_ of the four nanopores at 5.6*±*0.18 M, please see Figure S10 for the individual nanopore values. Previous calorimetry measurements ^70,71^ found the following guanidinium *K*_D_ values: 3.33 M to ubiquitin, 1.96 M to Cyt.*c*, 1.66 M to ribonuclease, and 1.44 M to lysozyme. The systematically lower values seen in those experiments can be explained by the unfolded state of the proteins, which provides more accessible sites for Gdm^+^ binding in comparison to the corresponding folded states.

Further analysis of the MD trajectories yields twenty residue type-specific *K*_D_ values (Figure 4g), which were insensitive to the polarity of the transmembrane bias (Figure S11). The magnitude of the computationally derived values is in general agreement with the results of previous measurements^72,73^ that found *K*_D_ to range from 40 to 90 mM for specific binding of Gdm^+^ to alanine, arginine, threonine, glutamic acid, aspartic acid, and tyrosine. The high binding affinity of Gdm^+^ to negatively charged amino acids explains the long residence time of Gdm^+^ at specific locations within the nanopore (peaks in Figure 4d). As such binding lasts considerably longer than the time it takes for a chloride ion to pass through the nanopore, the binding effectively acts as a permanent charge producing anion selectivity of the nanopore current.

### Ion selectivity and the electro-osmotic flow

Our simulations so far have shown that the presence of guanidinium induces moderate to strong anion selectivity in four biological nanopores. To determine how such selectivity translates into EOF, we characterized the displacement of water molecules through the nanopores in all our simulations. Figure 5a presents the results for the MspA nanopore, where guanidinium was found to produce the highest water flux despite rather modest anion selectivity. Thus, for 2.5 M KCl and at +200 mV bias, we find the water to flow through the MspA nanopore in the direction opposite the direction of the ionic current, conforming to the mild anion selectivity of the nanopore. Replacing K^+^ ions with Gdm^+^ increases the water flux by 3.6 times, despite the 46% overall reduction of the ionic current, Figure 3b. This observation is explained by the sustained anion current that reduces by only 1.3 times upon the cation replacement and almost complete abolishment of the cation current, which would carry the water in the direction of the ionic current, partially canceling out the flux caused by anion displacement. Similar magnitude EOF is measured for the 1.5 M GdmCl / 1.0 M KCl electrolyte mixture, which is consistent with the similar anion selectivity of the channel. For the latter condition, reversing the polarity of the transmembrane bias reversed the direction of the EOF and increased the EOF magnitude, consistent with the increased anion selectivity, Figure 3f.

**Figure 5:**
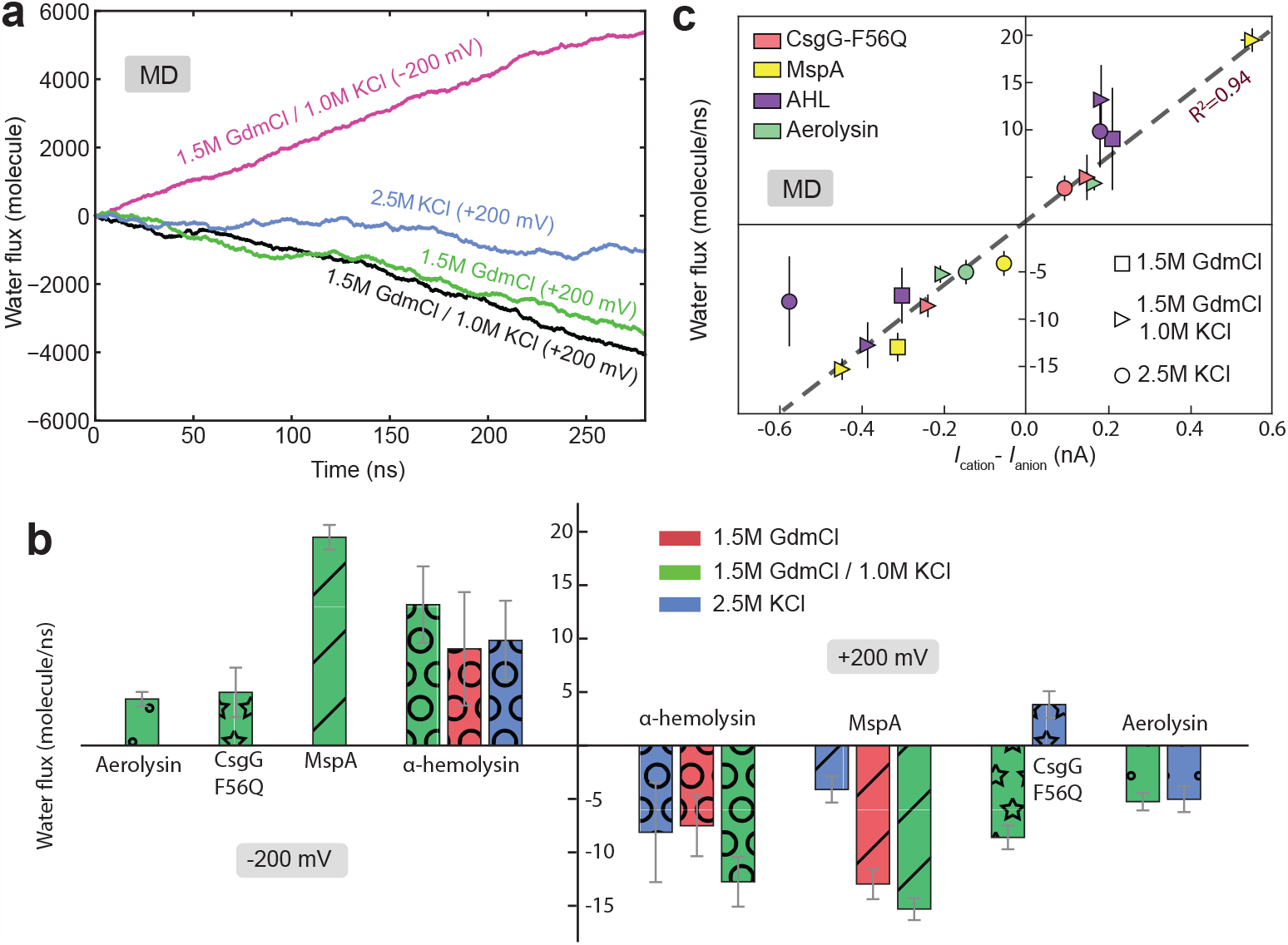
Electro-osmotic flow induced by guanidinium chloride in biological nanopores. **a**, Number of water molecules that permeated through the MspA constriction (residues 90, 91 and 93) *versus* simulation time. Negative values indicate transport in the negative *z* -axis direction, as defined in Figure 3a. **b**, Average water flux for each electrolyte condition in four biological nanopores. The average value and the standard error are calculated by splitting the trajectory into 30 ns fragments. **c**, Average water flux *versus* relative ionic current defined as the difference between the current carried by cationic and anionic species. The dashed line shows a linear fit to the data.

Figure 5b compares the average water flux for all four biological nanopores, the two transmembrane bias polarities and, for the MspA and *α*-hemolysin, the three electrolyte conditions. Please see Figure S1e-g for water flux traces of *α*-hemolysin simulations. Consistent with the observed ion selectivity, *α*-hemolysin and MspA systems were found to exhibit a strong electro-osmotic effect at either polarities of the transmembrane bias in the presence of Gdm^+^. Also consistent with the ion selectivity data (Figure 3f), the highest EOF for these two nanopores was observed in the case of the GdmCl / KCl electrolyte mixture.

For some biological nanopores, the presence of Gdm^+^ can reverse the direction of the EOF. Thus, we find the F56Q variant of the CsgG nanopore to exhibit cation selectivity in a pure KCl electrolyte.In the mixed KCl/GdmCl electrolyte, however, this nanopore exhibits anion selectivity, which reverses the direction of the EOF in comparison to the pure KCl system, Figure 5b and Figure S5d-f.

Under the same electrolyte and applied bias conditions, the magnitude of the EOF clearly depends on the nanopore type. Thus, aerolysin and MspA show the lowest and the highest water flux rate, respectively, (Figure 5a,b and Figure S4d-f). In order to establish a quantitative correlation between the water flux rate and the ionic current, we plot in Figure 5c the average water flux as a function of the difference between the average current carried by the cationic and anionic species—the relative ionic current, *I*_cation_ − *I*_anion_. The average water flux appears to linearly correlate with the relative ionic current quite well, taking into account the error in determining the water flux.^52^

### Sticky ions induce EOF in solid-state nanopores

Electroosmotic flow has been widely used in solid-state nanopore measurements in order to draw uncharged molecules such as proteins and other biopolymers.^14,74,75^ To investigate if the effect of guanidinium extends to solid-state nanopores, we built an all-atom model of a properly annealed SiO_2_ nanopore,^76,77^ a good computational representation of silicon nitride nanopores that are widely used in nanopore sensing.^13^ Two simulation systems were built each containing a 2.5 nm diameter nanopore in a 9.9 nm thick SiO_2_ membrane and either 2.5 M KCl or 1.5 M GmdCl / 1 M KCl electrolyte, the latter system is shown in Figure 6a. Subject to +200 mV, the nanopore current in the KCl system is found to be mildly cation selective (Figure 6b), which we attribute to the intrinsic cation selectivity of the SiO_2_ nanopore, whose surface is on average negatively charged.^78^ In the case of the electrolyte mixture containing guanidinium chloride, we find the nanopore current to be dominated by the flux of Cl^−^ ions (Figure 6c). Similar to biological pores, the presence of guanidinium chloride decreased the total ionic current (Figure 6d), while making the current anion selective. Accordingly, the presence of guanidinium chloride reversed the direction of the EOF from flowing along the direction of the ionic current to opposite to that (Figure 6e), increasing the EOF magnitude.

**Figure 6:**
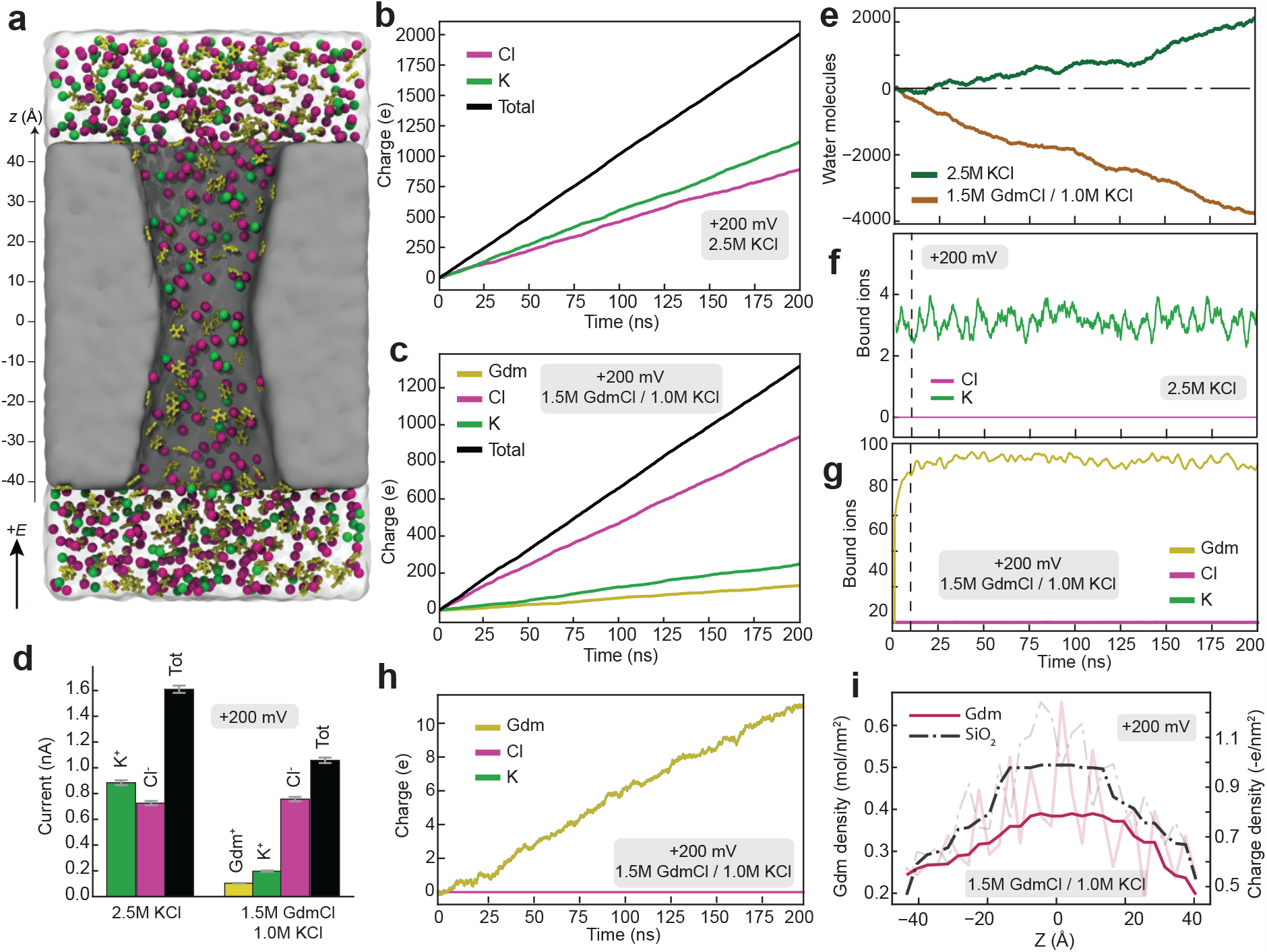
Guanidinium induces strong EOF in a solid-state nanopore. **a**, All-atom model of a SiO_2_ nanopore submerged in a mixture of 1.5 M GmdCl and 1.0 M KCl electrolytes. The nanopore is shown as a cut-away molecular surface. The semitransparent surface illustrates the volume occupied by the electrolyte. Gdm^+^, K^+^ and Cl^−^ ions are shown in yellow, green, and magenta, respectively. **b**,**c**, Total charge carried by ion species through a SiO_2_ nanopore over the course of MD simulations performed at +200 mV and 2.5 M KCl (panel b) or 1.5 M GmdCl / 1 M KCl (panel c) electrolyte. **d**, Average current carried by each ion species. **e**, Number of water molecules that permeated through the constriction (−30 *< z <* 30 Å) of the SiO_2_ nanopore.Negative values indicate transport in the negative *z* -axis direction, as defined in panel a. **f**,**g**, Number of ions bound to the inner surface of the nanopore (excluding the top and bottom surfaces of the membrane) for the 2.5M KCl (f) and the 1.5 M GdmCl / 1.0 M KCl (g) simulation. An ion is considered bound if any of its atoms are located within 3 Å of any atom of the SiO_2_ membrane. **h**, Total charge carried by surface bound ions as a function of simulation time. **i**, Surface density of Gdm^+^ bound to the SiO_2_ nanopore (solid lines, left axis) and the charge density of the nanopore surface (dashed line, right axis). The densities were calculated in 3 Å bins along the *z* -axis and smoothed using a median filter with a sampling window of 13 Å.

To elucidate the molecular causes of the EOF reversal, we computed the number of ions bound to the SiO_2_ surface for the KCl and mixed electrolyte simulations. In the case of the KCl electrolyte, no Cl^−^ were found to bind to the nanopore surface while less than 4 K^+^ ions where found, on average, to reside within 3 Å of the surface (Figure 6f). However, for the mixed electrolyte, the number of Gdm^+^ ions near the nanopore surface was found to increase with simulation time, reaching a steady-state of 91*±*2 Gdm^+^ bound after about 30 ns of the simulation time (Figure 6g). Just like for biological nanopores, the binding is not permanent and Gdm^+^ ions are observed to bind to, move along, and unbind from the nanopore surface. Although the surface-bound Gdm^+^ can move along the surface, the contribution of such surface-bound current is relatively small: less then 12 ions were found to pass through the nanopore constriction, on average, over the course of the MD simulation (Figure 6h), while the total number of Gdm^+^ ions transported through the nanopore over the same time interval is 131. The density of the Gdm^+^ ions bound to the SiO_2_ surface is found to gradually increase from the nanopore entrance to the nanopore constriction and correlate with the local charge of the nanopore surface (Figure 6i). Thus, the presence of guanidinium ions can have a pronounced effect on the magnitude and direction of the EOF in a solid-state nanopore *via* the same surface-binding mechanism as in biological nanopores.

## Conclusions

We have described a general method for inducing a strong electro-osmotic effect in biological and solid-state nanopores. We have showed that the addition of high concentrations of guanidinium chloride to the solution used for nanopore translocation experiments makes the nanopore anion selective. The effect is found to originate from transient sticking of guanidinium ions to the inner surface of the nanopores, which renders the surface positively charged. The local electroneutrality condition requires such positive charge at the nanopore surface to be compensated by an equivalent charge of mobile anions residing within the nanopore volume. The presence of such compensating ions gives the solution inside the nanopore a net electrical charge. Subject to an external electric field, the charged solution moves in the direction prescribed by he applied field, generating a strong electro-osmotic effect.

Unlike the methods previously described to induce the ion selectivity and the electroosmotic flow in biological nanopores, our approach does not require any modifications to the nanopore shape, introduction of local charge through point mutations, or the use of asymmetric electrolyte conditions. Our approach can also reverse and/or enhance the ion selectivity and the electro-osmotic effect in solid-state nanopores, which in previous work required either chemical treatment of the nanopore surface or the use of additional electrodes to locally influence the nanopore electrostatics. Furthermore, the sticky-ion action is fully reversible and should remain effective under a wide range of pH conditions, as long as the nanopore retains its structure. We expect the results of our work to find applications in a variety of technological areas, in particular in nanopore sensing and sequencing, providing an easy-to-follow recipe for transporting heterogeneously charged and uncharged (bio)molecules unidirectionally through nanoscale passages. Finally, we note that the transient binding principle is not limited to guanidinium ions and can be reproduced using other small molecules as long as they can exhibit moderate affinity to the nanopore surface and carry electrical charge, with one such potential candidate being sodium dodecyl sulfate. ^79^

## Acknowledgement

This work was supported by the grant from the National Institutes of Health (R01-HG012553 to B.M., L.Y., M.W., and A.A). The supercomputer time was provided through the ACCESS allocation MCA05S028 and the Leadership Resource Allocation MCB20012 on Frontera of the Texas Advanced Computing Center.

## Supporting Information Available

This information is available free of charge via the Internet at http://pubs.acs.org.

MD methods; nanopore experiments; comparison of simulated and experimental open-pore ionic current through *α*-hemolysin; experimental and simulated bulk conductivity of three electrolyte solutions; activity coefficients for KCl and GdmCl electrolytes; MD simulation results for *α*-hemolysin, MspA, aerolysin, and CsgG systems; experimentally measured current-voltage dependence of *α*-hemolysin; simulated local ion concentration in the transmembrane pore of *α*-hemolysin; analysis of Gdm^+^ binding to the inner surface of *α*-hemolysin; number of Gdm^+^ ions bound to the inner surface of the four transmembrane pores as a function of simulation time; amino acid specific interaction of Gdm^+^ with biological nanopores; overall binding affinity of Gdm^+^ to biological nanopores; binding affinity of Gdm^+^ to amino acids of biological nanopores.

---

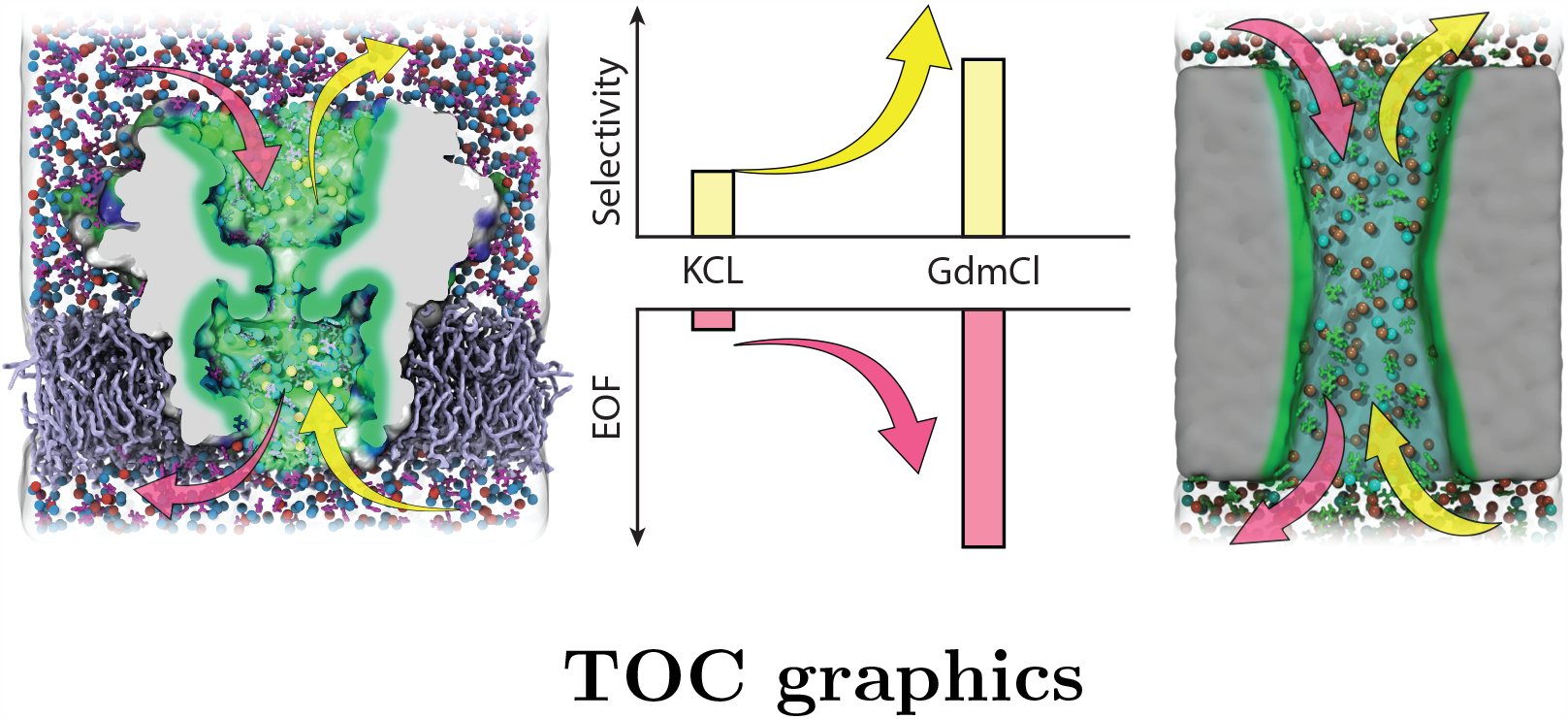

