## Supporting Information for "Electro-Osmotic Flow Generation via a Sticky Ion Action"

### Material and Methods

#### General MD Methods

All MD simulations were performed using NAMD,[1] the CHARMM36 parameters for protein and lipid molecules,[2–4] the CUFIX corrections to charge-charge interactions,[5,6] and the TIP3P model of water.[7] Unless specified otherwise, a 2 fs integration time step was used along with the SETTLE[8] and RATTLE[9] algorithms for restraining the covalent bonds of hydrogens in water or protein and lipids, respectively. All simulations employed periodic boundary conditions in three dimensions. The electrostatic interactions were calculated by applying the Particle mesh Ewald (PME) algorithm over a 1 Å spaced grid.[10] The van der Waals forces were evaluated using a 10–12 Å smooth cutoff. Local interactions were evaluated every time step and full electrostatics every third time step. In the constant number of particles, volume and temperature (NVT) simulations, Langevin thermostat[11] was applied to all the heavy atoms of the membrane with a damping coefficient of 1 $ps^{-1}$ to keep the system's temperature at 298 K. Constant pressure (NPT) simulations used the Nosé-Hoover Langevin piston pressure control[12] to maintain the pressure at 1 bar. Energy minimization was performed using a conjugate gradient method. During the NPT equilibration, the area of the lipid bilayer was kept constant while the system was allowed to change its size along the bilayer normal. All production simulations were carried out in the NVT ensemble under a constant external electric field applied normal to the membrane, producing a ±200 mV transmembrane voltage. All $C_\alpha$ atoms of the protein nanopores were harmonically restrained to their crystallographic coordinates; the spring constant of each restraint was 0.1 kcal $mol^{-1}Å^{-2}$. Visualization and analysis were performed using VMD.[13]

#### All-atom models of biological nanopores

Our all-atom models of $\alpha$-hemolysin nanopores were constructed and equilibrated as described previously.[14] Briefly, each model included one $\alpha$-hemolysin protein merged with a

$15 \times 15$ nm$^2$ patch of a POPC bilayer and solvated with a rectangular volume of water. Ions were first introduced to neutralize the system, upon which $Gdm^+$, $K^+$, and $Cl^-$ ions were added by replacing an equivalent number of water molecules and removing additional water molecules located within 2.4 Å of each ion. Three $\alpha$-hemolysin systems were constructed differing by their electrolyte composition, namely 1.5 M GdmCl, 1.5 M GdmCl / 1.0 M KCl, and 2.5 M KCl. The final systems measured $15 \times 15 \times 18$ nm$^3$ in volume and contained approximately 300,000 atoms. The assembled systems were energy-minimized for 5,000 steps, followed by a 2.5 ns equilibration simulation in which the protein backbone and the lipid head groups were restrained. This step was followed by a 25 ns simulation in the NPT (constant number of particles, pressure and temperature) ensemble using the Nose-Hoover Langevin piston pressure control.

Our all-atom models of M1-NNN MspA were constructed as described previously.[15] Briefly, the crystallographic structure of the wild-type MspA (PDB ID 1UUN)[16] was used to build the M1-NNN-model of MspA by reverting all R96 residues to alanines and imposing the D90N/D91N/D93N mutations. After aligning the primary principal axis of the protein with the $z$-axis, the protein structure was merged with a $12 \times 12$ nm$^2$ patch of pre-equilibrated 1,2-diphytanoyl-sn-glycero3-phosphocholine (DPhPC) bilayer. All DPhPC molecules that overlapped with the atoms of MspA were removed. The protein–lipid assembly was immersed in a rectangular volume of water, producing a system containing approximately 164,000 atoms. $Gdm^+$, $K^+$, and $Cl^-$ ions were added at random positions corresponding to target ionic concentrations and electrically neutralizing the system. Similar to the $\alpha$-hemolysin systems, three MspA systems were built differing by the electrolyte composition. Upon assembly, each system was energy-minimized for 2,000 steps and equilibrated for 2 ns having the protein backbone and the lipid head groups harmonically restrained with spring constants of 1.0 kcal mol$^{-1}$Å$^{-2}$. Following that, the systems were equilibrated for 23 ns with only the backbone of the protein restrained with spring constants set to 0.1 kcal mol$^{-1}$Å$^{-2}$.

Our all-atom model of aerolysin nanopore was built starting from a previously described,[17] pre-equilibrated (20 ns) model containing a wild-type aerolysin protein embedded in a $19 \times 19$ nm$^2$ DPhPC bilayer. Water and ions were added to each system using the 'Solvate' and 'Autoionize' plugins of VMD.[13] The number of $Gdm^+$, $K^+$, and $Cl^-$ ions was adjusted to produce an electrically neutral system containing a 1.5 M GdmCL / 1.0 M KCl solution. Upon assembly, the system was energy-minimized for 5,000 steps and then equilibrated for 25 ns in a constant number of atoms (503,411), pressure (1 bar), and temperature (298 K) ensemble. During this process, all $C_\alpha$ atoms of the protein were restrained to their initial coordinates.

Our all-atom model of a CsgG nanopore was constructed using the crystal structure of the WT-CsgG pore (PDB ID 4UV3).[18] The missing atoms in the crystallographic structure and the F56Q mutation were incorporated in the model using the psfgen package of VMD [11]. The CsgG pore axis was aligned with the $z$-axis of our coordinate system. The protein was then merged with a $14 \times 14$ nm$^2$ pre-equilibrated patch of a POPC bilayer. Lipids that overlapped with the protein were removed. The protein-lipid complex was then solvated in a rectangular volume of water. Next, $Gdm^+$, $K^+$, and $Cl^-$ ions were added to the system to produce a 1.5 M GdmCl / 1.0 M KCl electrolyte solution. The final system measured $14 \times 14 \times 16.5$ nm$^3$ in volume and contained approximately 305,000 atoms. The system underwent 5,000 steps of energy minimization and NPT equilibration having all $C_\alpha$ atoms of the protein harmonically restrained to their initial coordinates. For the first 10 ns, the spring constant of each restraining potential was 1.0 kcal mol$^{-1}$Å$^{-2}$, which was then reduced to 0.5 kcal mol$^{-1}$Å$^{-2}$ and then to 0.1 kcal mol$^{-1}$Å$^{-2}$ in 10 ns intervals.

#### All-atom models of solid-state nanopores

Our model of $SiO_2$ nanopore was built by generating a $4.5 \times 10$ nm$^2$ hexagonal patch (4.5 nm on side and 10 nm in height) of $SiO_2$ using the Inorganic Builder plugin of VMD.[13] Force field parameters describing interactions of $SiO_2$ atoms were taken from Ref. 19. A double-cone nanopore was then generated in the silica slab and the system was annealed as described in Ref. 20. Covalent bonds were defined to pairs of adjacent $SiO_2$ conforming to hexagonal periodic boundary conditions in the $xy$ plane. Next, the membrane was solvated with a 4 nm-thick volume of water on each side. Two $SiO_2$ systems were built containing either a 2.5 M KCl electrolyte or a 1.5 M GdmCL / 1.0 M KCl electrolyte mixture. Each system underwent energy minimization followed by 30 ns equilibrium in which the atoms of $SiO_2$ were harmonically restrained (with the force constant of 20 kcal mol$^{-1}$Å$^{-2}$) to their coordinates obtained at the end of the annealing procedure. All $SiO_2$ simulations were carried out using a time step of 1 fs.

#### Analysis of the MD trajectories

The ionic currents were calculated following a previously described method.[14] The concentration profile and guanidinium binding analyses were performed using Python and Tcl scripts developed in-house. The electrostatic potential was determined using a method described previously,[14] which was implemented in the PMEpot plugin of VMD. The method averages the instantaneous distributions of the electrostatic potential over the respective MD trajectory.

The dissociation constant, $K_D$, between guanidinium ions and a protein was measured assuming the protein to be a macromolecule containing identical independent binding sites. Under this assumption and in equilibrium, the dissociation coefficient, $K_D$, is obtained as:[21]

$$\frac{[\mathrm{Gdm}]}{K_D + [\mathrm{Gdm}]} = \frac{[\mathrm{Gdm}]_{\mathrm{bound}}}{n[\mathrm{Protein}]} \tag{1}$$

where [Gdm] and $[\mathrm{Gdm}]_{\mathrm{bound}}$ represent the concentration of the guanidinium in the solution and bound to the protein, respectively. Above, [protein] denotes the concentration of the protein and $n$ is the number of the identical independent binding site on the protein. For the non-specific ligand-binding site analysis, all residues on the surface of the proteins were assumed as identical binding pockets. However, for site-specific Gdm binding, the number of binding pockets was equal to the total number of the amino acid of interest exposed to the electrolyte solution.

#### Nanopore experiments

The 500 $\mu$m-thick silicon chip with 20 $\mu$m-thick SU-8 photoresist on the top layer was mounted on our customized fluidic cell, sealing properly, to separate *cis* and *trans* chambers. A 100 $\mu$m wedge-on-pillar shaped aperture was designed as a biomimetic bilayer membrane supporter on the SU-8 layer. Both sides of the aperture were pretreated with 4 mg/ml poly(1,2-butadiene)-b-poly(ethylene oxide) (PBD11–PEO8) block-copolymer (Polymer Source) dissolved in hexane. After complete evaporation of the solvent, the aperture was coated with a dry and thin polymer layer. The *cis* and *trans* chambers were filled with electrolyte, and a pair of Ag/AgCl electrodes that connected to an Axon 200B patch-clamp amplifier were inserted in the electrolyte. A polymer membrane was formed across the aperture by applying 4 mg/ml polymer (dissolved in decane) solution on it. After thinning the membrane to bilayer by bubbling using a pipet, 0.5 $\mu$l of 0.5 $\mu$g/ml $\alpha$-hemolysin (Sigma-Aldrich) was added to the *cis* chamber, and an ion conductance jump marked single pore insertion. Current signals were low-pass filtered at 10 kHz using the Axopatch setting and digitized at 16-bits and 250 kHz sampling rate using a National Instruments Data Acquisition card and custom LabVIEW-based software that records and saves all raw current data and acquisition settings.

#### Ion selectivity analysis of experimental measurements

Since we used two wire Ag/AgCl electrodes whose potential at each chamber depends on the $Cl^-$ concentration in the electrolyte, the experimentally applied voltage, $V_\mathrm{m}$, was first corrected by the Nernst potential, $V_\mathrm{N}$, such that

$$V_\mathrm{c} = V_\mathrm{m} - V_\mathrm{N} \tag{2}$$

and Nernst potential $V_N$ was calculated as

$$V_{\mathrm{N}} = \frac{RT}{nF} \ln \frac{[a_{\mathrm{Cl^-}}]_{cis}}{[a_{\mathrm{Cl^-}}]_{trans}}, \tag{3}$$

where $R$ is the ideal gas constant, $T$ is the absolute temperature, $F$ is the Faraday's constant, $n = 1$ is the number of electrons in the electrochemical reaction, and $[a_{\mathrm{Cl^-}}]$ is $\mathrm{Cl^-}$ activity in the corresponding electrolyte chamber, which equals to the product of the activity coefficient $\gamma$ and the ion concentration:

$$[a_{\mathrm{Cl^-}}] = \gamma[\mathrm{Cl^-}]. \tag{4}$$

Reversal potential, $V_r$, measurements probe the electrical potential difference that is created across an ion-selective membrane, or a channel placed between two solutions that differ in composition and/or concentration of salts. Thus, with all $I$-$V$ curves presented as $I$ *versus* $V_{\mathrm{c}}$, the reversal potential $V_{\mathrm{r}}$ is the value of $V_{\mathrm{c}}$ where $I = 0$ (see Figure 1e, and SI Figure 2 for uncorrected current-voltage curves). Reversal potential probes the relative permeance of each ion through the pore, and the general form of $V_{\mathrm{r}}$ when a channel is in contact with a salt concentration gradient reads

$$V_{\mathrm{r}} = \frac{RT}{F} \ln \frac{\sum_i^n P_{M_i^+}[a_{M_i^+}]_{cis} + \sum_j^m P_{A_j^-}[a_{A_j^-}]_{trans}}{\sum_i^n P_{M_i^+}[a_{M_i^+}]_{trans} + \sum_j^m P_{A_j^-}[a_{A_j^-}]_{cis}}, \tag{5}$$

where $P$ is the ion permeability and $a$ is the ion activity. For the case of 1 M $[\mathrm{KCl}]_{cis}$ : 0.1 M $[\mathrm{KCl}]_{trans}$ (solid triangle markers in Figure 1d), Equation 5 becomes

$$V_{\mathrm{r}} = \frac{RT}{F} \ln \frac{P_{\mathrm{K}}[a_{\mathrm{K^+}}]_{cis} + P_{\mathrm{Cl}}[a_{\mathrm{Cl^-}}]_{trans}}{P_{\mathrm{K}}[a_{\mathrm{K^+}}]_{trans} + P_{\mathrm{Cl}}[a_{\mathrm{Cl^-}}]_{cis}}. \tag{6}$$

Dividing the numerator and the denominator in the logarithmic term by $P_{\mathrm{Cl}}$, we obtain

$$V_{\mathrm{r}} = \frac{RT}{F} \ln \frac{\frac{P_{\mathrm{K}}}{P_{\mathrm{Cl}}}[a_{\mathrm{K^+}}]_{cis} + [a_{\mathrm{Cl^-}}]_{trans}}{\frac{P_{\mathrm{K}}}{P_{\mathrm{Cl}}}[a_{\mathrm{K^+}}]_{trans} + [a_{\mathrm{Cl^-}}]_{cis}}. \tag{7}$$

Using activity coefficients for 1 M and 0.1 M KCl of 0.604 and 0.768, respectively[22] (see SI Table 3), Equation 7 becomes

$$V_{\mathrm{r}} = \frac{RT}{F} \ln \frac{0.604\frac{P_{\mathrm{K}}}{P_{\mathrm{Cl}}} + 0.0768}{0.0768\frac{P_{\mathrm{K}}}{P_{\mathrm{Cl}}} + 0.604}. \tag{8}$$

To calculate the ion selectivity for $\alpha$-hemolysin in contact with the GdmCl concentration gradient, we used known $\gamma$ values for 1 M and 0.1 M GdmCl of 0.497 and 0.768, respectively,[23] and modified Equation 5 as follows

$$V_{\mathrm{r}} = \frac{RT}{F} \ln \frac{P_{\mathrm{G}}[a_{\mathrm{Gdm}^+}]_{cis} + P_{\mathrm{Cl}}[a_{\mathrm{Cl}^-}]_{trans}}{P_{\mathrm{G}}[a_{\mathrm{Gdm}^+}]_{trans} + P_{\mathrm{Cl}}[a_{\mathrm{Cl}^-}]_{cis}}. \tag{9}$$

**SI Table 1. Simulated and experimental open pore ionic current through $\alpha$-hemolysin.** The data in the $I^*_{\mathrm{MD}}$ column specifies the simulated MD currents multiplied by the ratio of the experimental and simulated bulk electrolyte conductivities, Table S2. The last two columns compare the simulated and experimental ionic current rectification ratio $\left|\frac{I_{+200}}{I_{-200}}\right|$ for the three electrolyte conditions. The experimental data are from Ref. 24.

| Electrolyte compostion | $V$ (mV) | $I_{\mathrm{MD}}$ (nA) | $I^*_{\mathrm{MD}}$ (nA) | $I_{\mathrm{Exp}}$ (nA) | $\left\|\frac{I_{+200}}{I_{-200}}\right\|_{\mathrm{MD}}$ | $\left\|\frac{I_{+200}}{I_{-200}}\right\|_{\mathrm{Exp}}$ |
|---|---|---|---|---|---|---|
| 1.5 M GdmCl | +200 | +0.327 | +0.253 | +0.212 | 1.27 | 1.32 |
| | −200 | −0.257 | −0.199 | −0.160 | | |
| 1.5 M GdmCl/1.0M KCl | +200 | +0.471 | +0.461 | +0.332 | 1.10 | 1.06 |
| | −200 | −0.426 | −0.426 | −0.313 | | |
| 2.5 M Cl | +200 | +1.084 | +1.120 | +0.499 | 1.23 | 1.35 |
| | −200 | −0.880 | −0.909 | −0.369 | | |

**SI Table 2. Experimental and simulated bulk conductivity of three electrolyte solutions.** The simulated bulk conductivity was calculated by first simulating the ionic current through a 10×10×14 $nm^3$ volume of each electrolyte under six voltages in the ±300 mV range. The bulk conductivity $\sigma = \frac{I}{V}\frac{L}{A}$, where $L$ and $A$ are the length and the cross section area of the simulation system, respectively, $V$ is the voltage applied over length $L$ and $I$ is the ionic current passing through cross section $A$.

| Electrolyte compostion | Method | $\sigma(S/m)$ |
|---|---|---|
| 1.5 M GdmCl | MD | 14.54 |
| | Experiment | 11.28 (pH 4.8) |
| | | 11.32 (pH 7.5) |
| 1.5 M GdmCl / 1.0 M KCl | MD | 19.73 |
| | Experiment | 19.34 (pH 4.8) |
| | | 19.52 (pH 7.5) |
| 2.5 M KCl | MD | 23.85 |
| | Experiment | 24.65 (pH 7.5) |

**SI Table 3.** Activity coefficients for KCl[22] and GdmCl[23] electrolytes at 298 K.

| **Salt** | $\gamma$ |
|---|---|
| 0.1 M KCl | 0.768 |
| 1 M KCl | 0.604 |
| 0.1 M GdmCl | 0.768 |
| 1 M GdmCl | 0.497 |

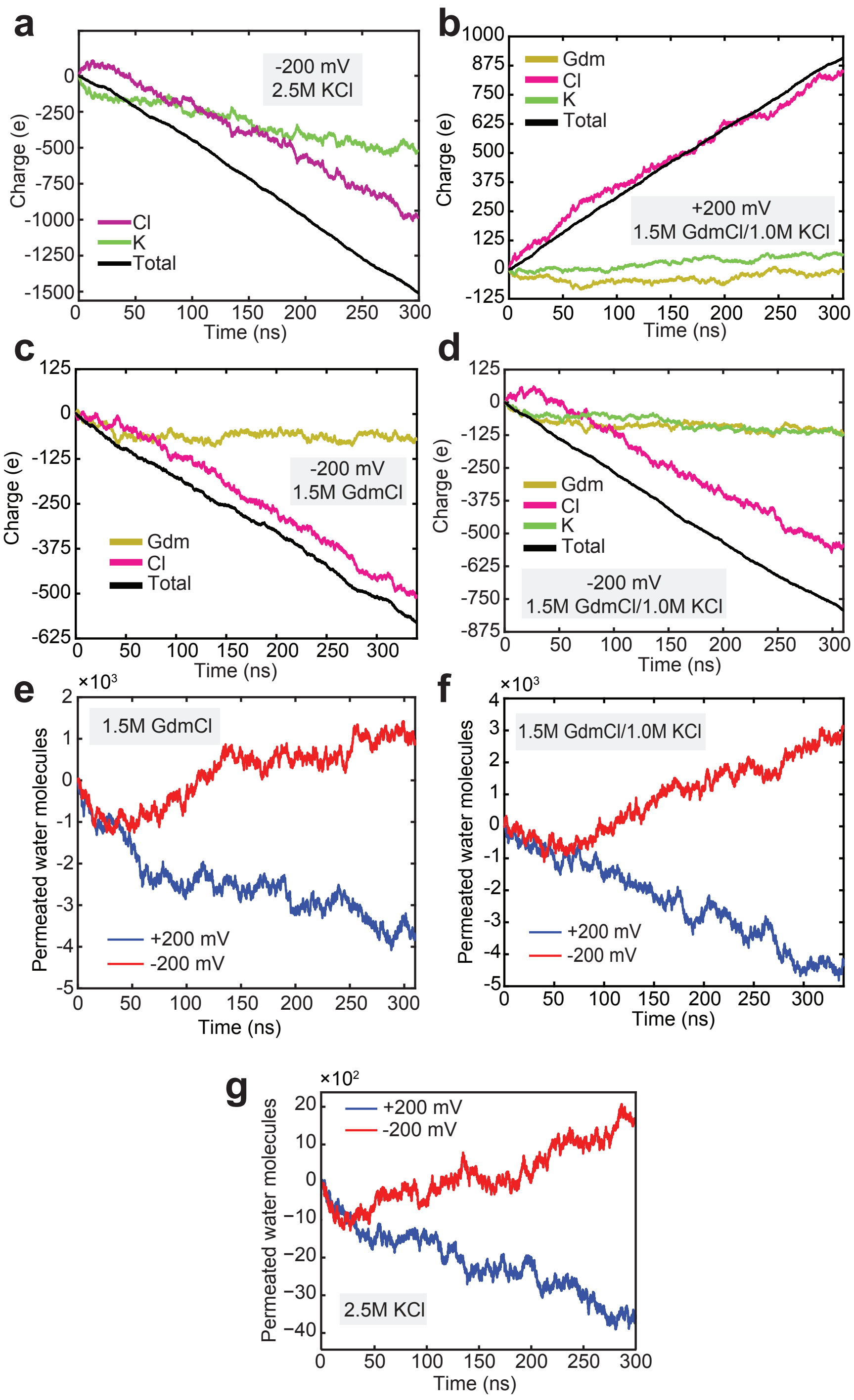

**SI Figure 1. MD simulations of $\alpha$-hemolysin systems. a-d,** Total charge carried by ion species through the transmembrane pore of $\alpha$-hemolysin as a function of simulation time. **e-g,** Number of water molecules permeated through the $\alpha$-hemolysin constriction (defined by residues 111, 113, and 147) as a function of simulation time. The electrolyte and applied bias conditions are specified in each panel.

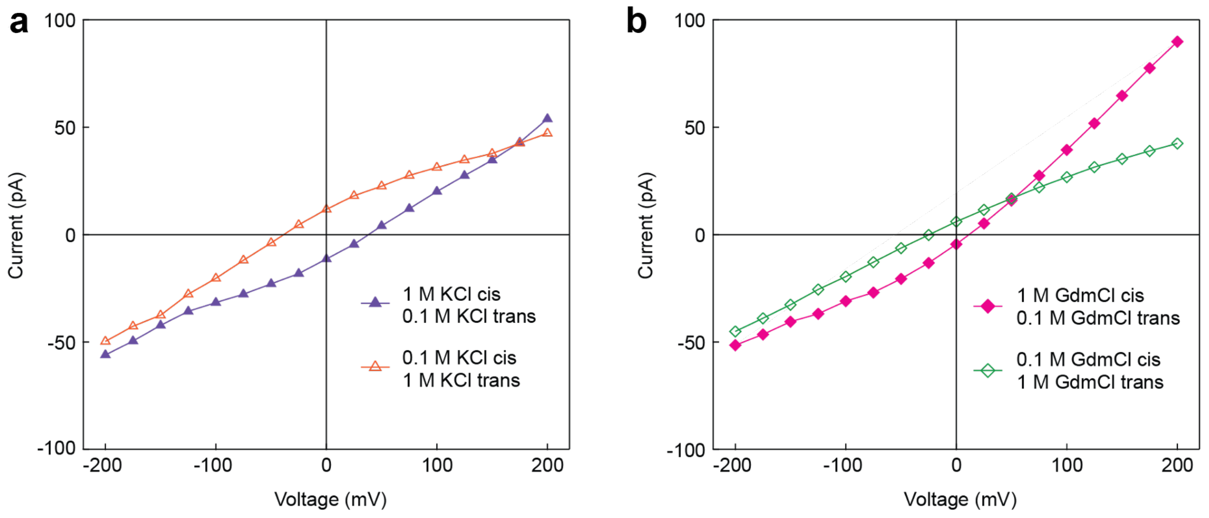

**SI Figure 2. Experimentally measured current-voltage dependence of $\alpha$-hemolysin.** The experimental conditions are specified in each panel.All buffers contained 10 mM Tris at pH 7.5. These row I-V curves do not account for the Nernst potential at the electrodes.

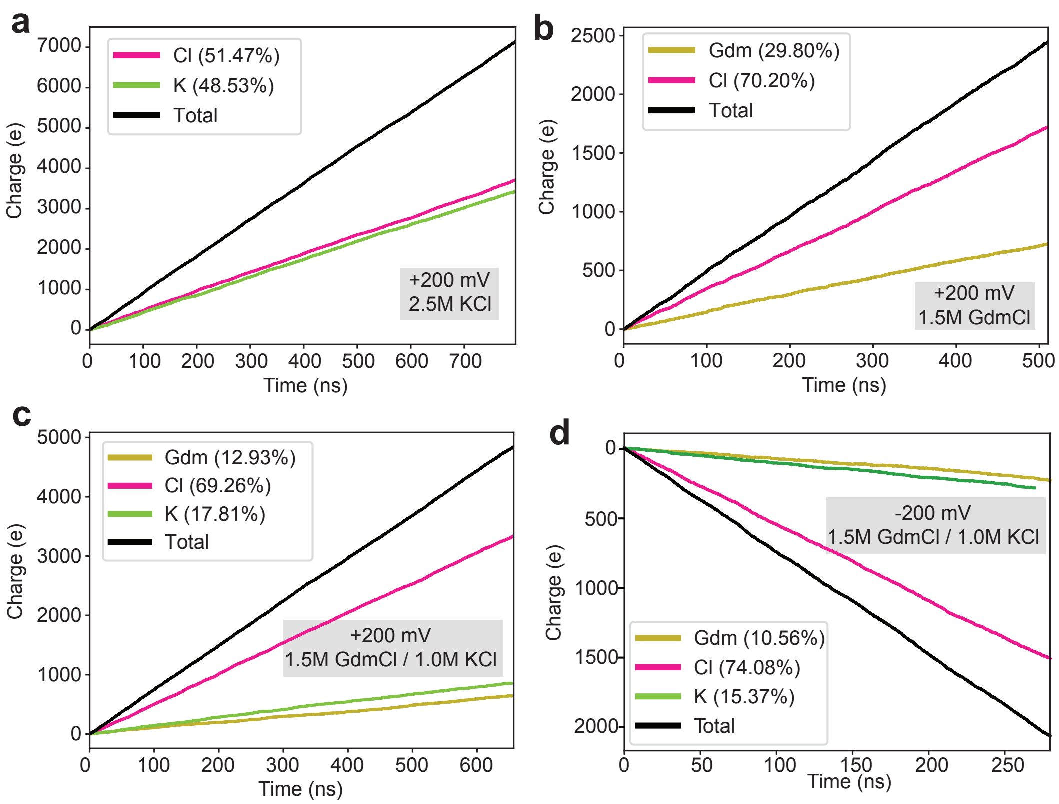

**SI Figure 3. MD simulations of MspA systems.** Total charge carried by ion species through the transmembrane pore of MspA as a function of simulation time. The electrolyte and applied bias conditions are specified in each panel. The EOF data are presented in main text Figure 5a.

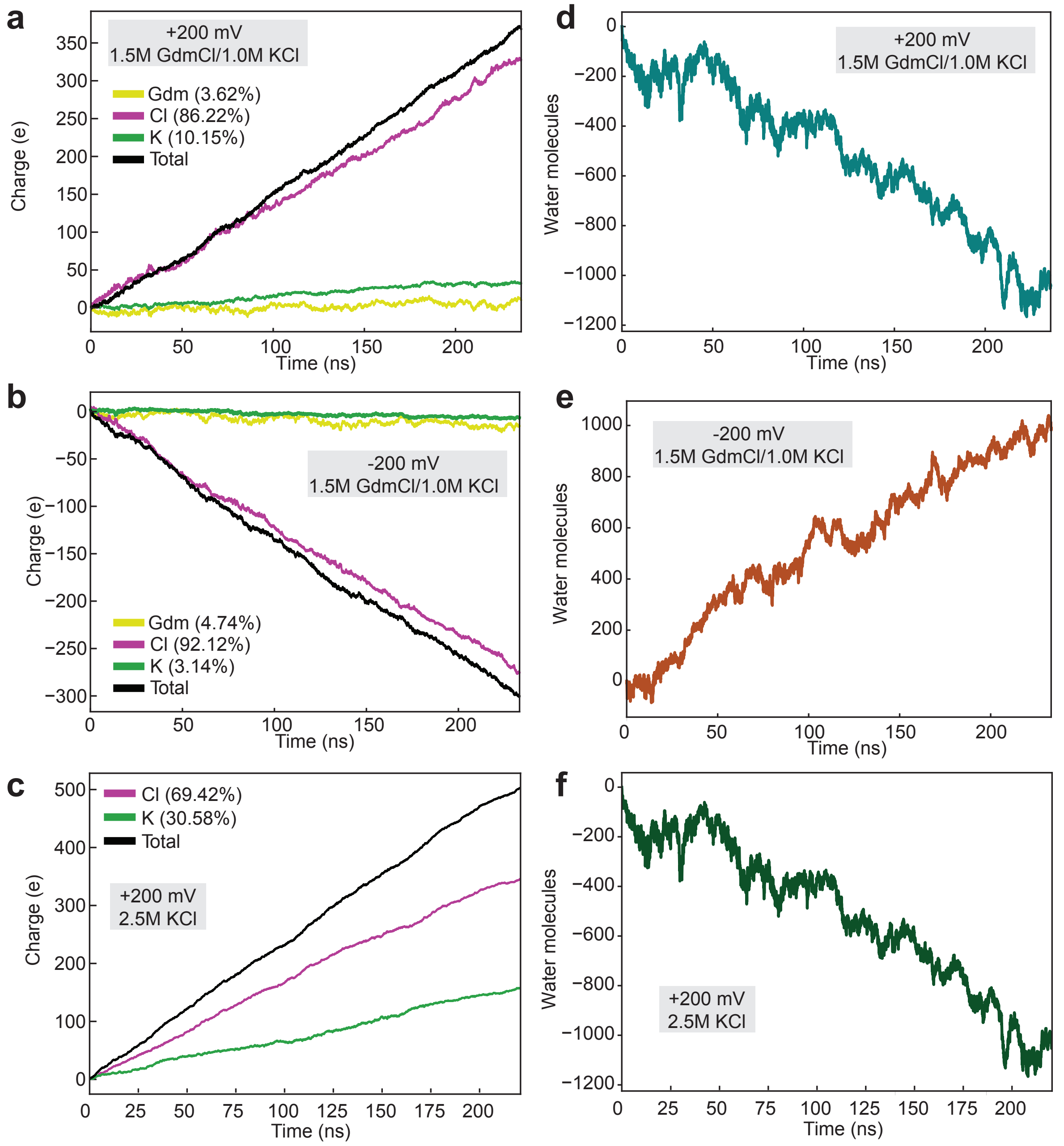

**SI Figure 4. MD simulations of aerolysin systems. a-c,** Total charge carried by ion species through the transmembrane pore of aerolysin as a function of simulation time. **d-f,** Number of water molecules permeated through the aerolysin constriction (defined by residues 220 and 238) as a function of simulation time. The electrolyte and applied bias conditions are specified in each panel.

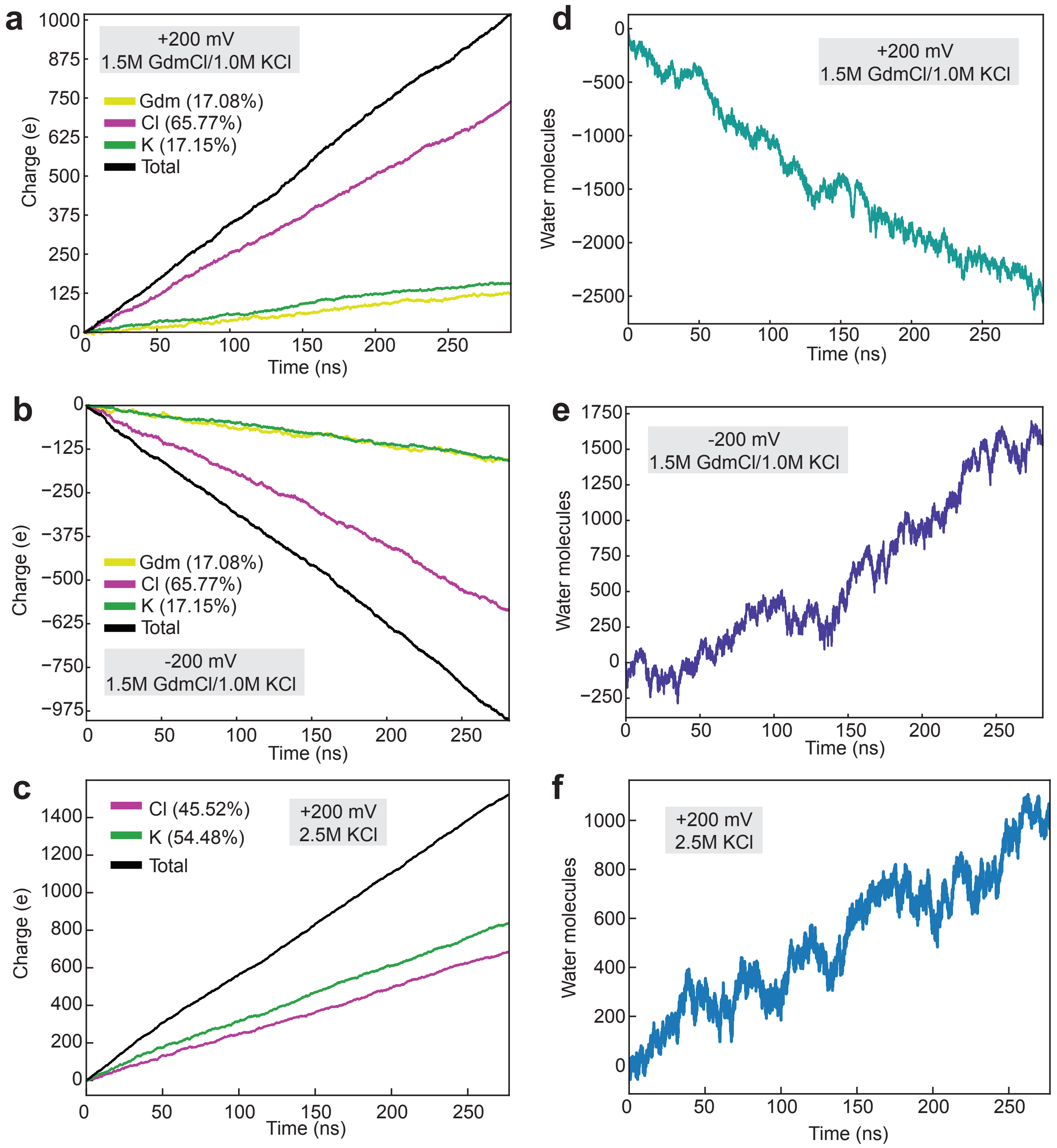

**SI Figure 5. MD simulations of CsgG systems. a-c,** Total charge carried by ion species through the transmembrane pore of aerolysin as a function of simulation time. **d-f,** Number of water molecules permeated through the CsgG constriction (defined by residues 51, 55, and 56) as a function of simulation time. The electrolyte and applied bias conditions are specified in each panel.

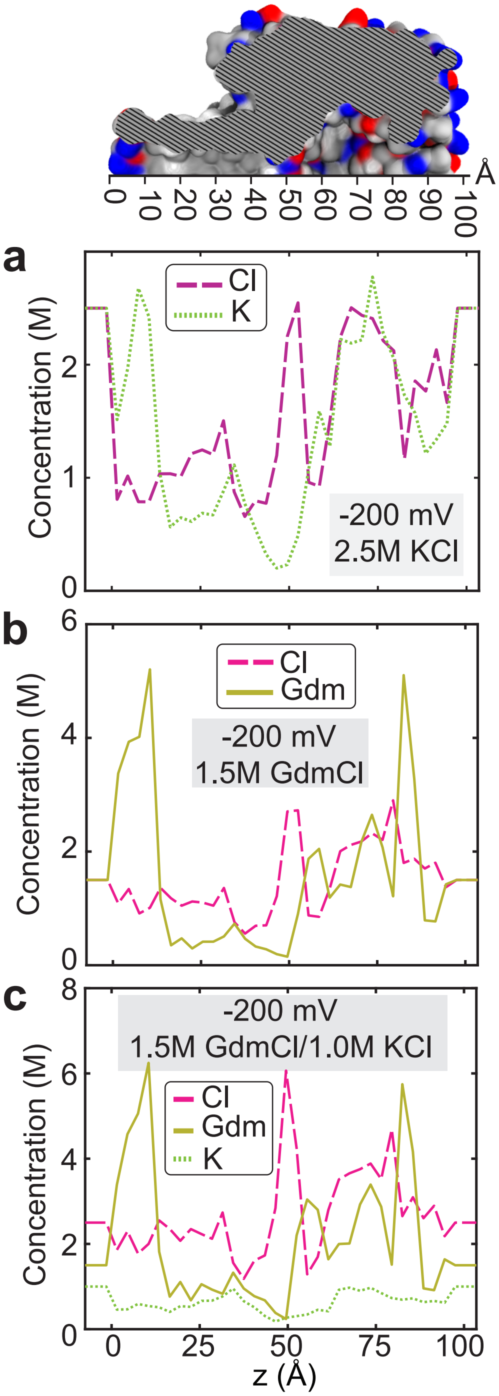

**SI Figure 6. Simulated local ion concentration in the transmembrane pore of $\alpha$-hemolysin.** The local concentrations were computed by averaging instantaneous values over the respective MD trajectories using 3 Å bins along the nanopore axis (the $z$-axis). The $z$-axis is defined in Figure 1A. The electrolyte and applied bias conditions are specified in each panel.

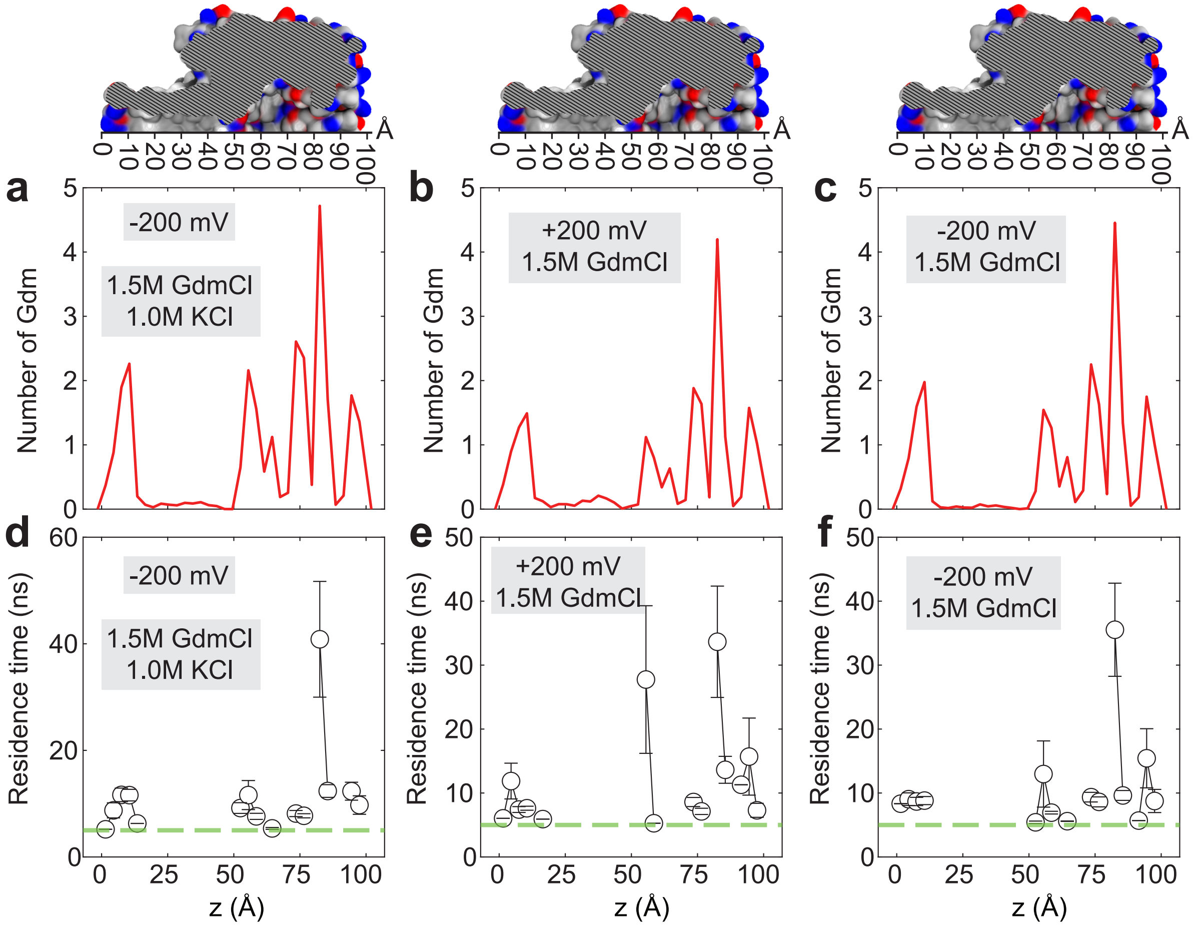

**SI Figure 7. Local binding of Gdm$^+$ to the inner surface of $\alpha$-hemolysin. a–c,** Average number of Gdm ions located within 3 Å of the inner surface of the $\alpha$-hemolysin nanopore. **d–f,** Average residence time of the Gdm$^+$ ions bound to the inner surface of the $\alpha$-hemolysin nanopore. The $z$-axis is defined in Figure 1A. Contacts between Gdm$^+$ and the nanopore surface lasting less than 5 ns (dashed green line) were discarded from the residence time analysis. The electrolyte and applied bias conditions are specified in each panel.

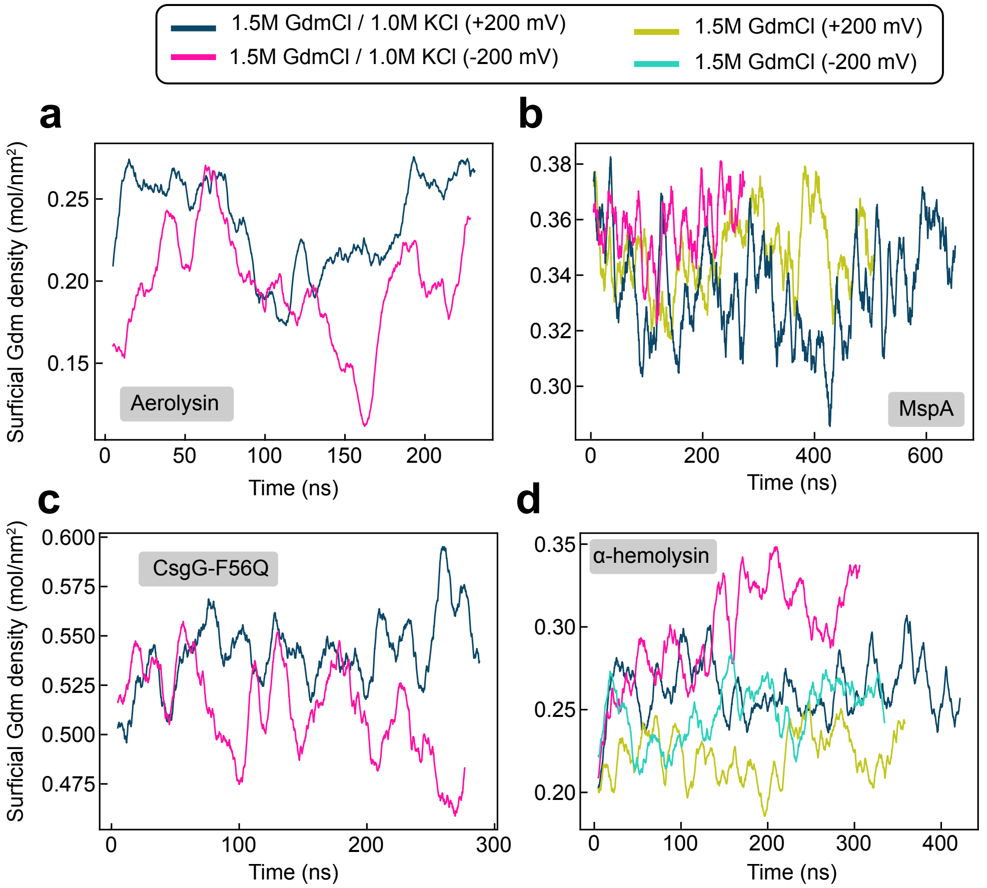

**SI Figure 8. Number of $Gdm^+$ ions bound to the inner surface of the four transmembrane pores as a function of simulation time.** The nanopore type is specifies in each panel. The electrolyte and applied bias conditions are specified above the graphs. Each curve shows a 5-ns window running average of the instantaneous data sampled every 10 ns.

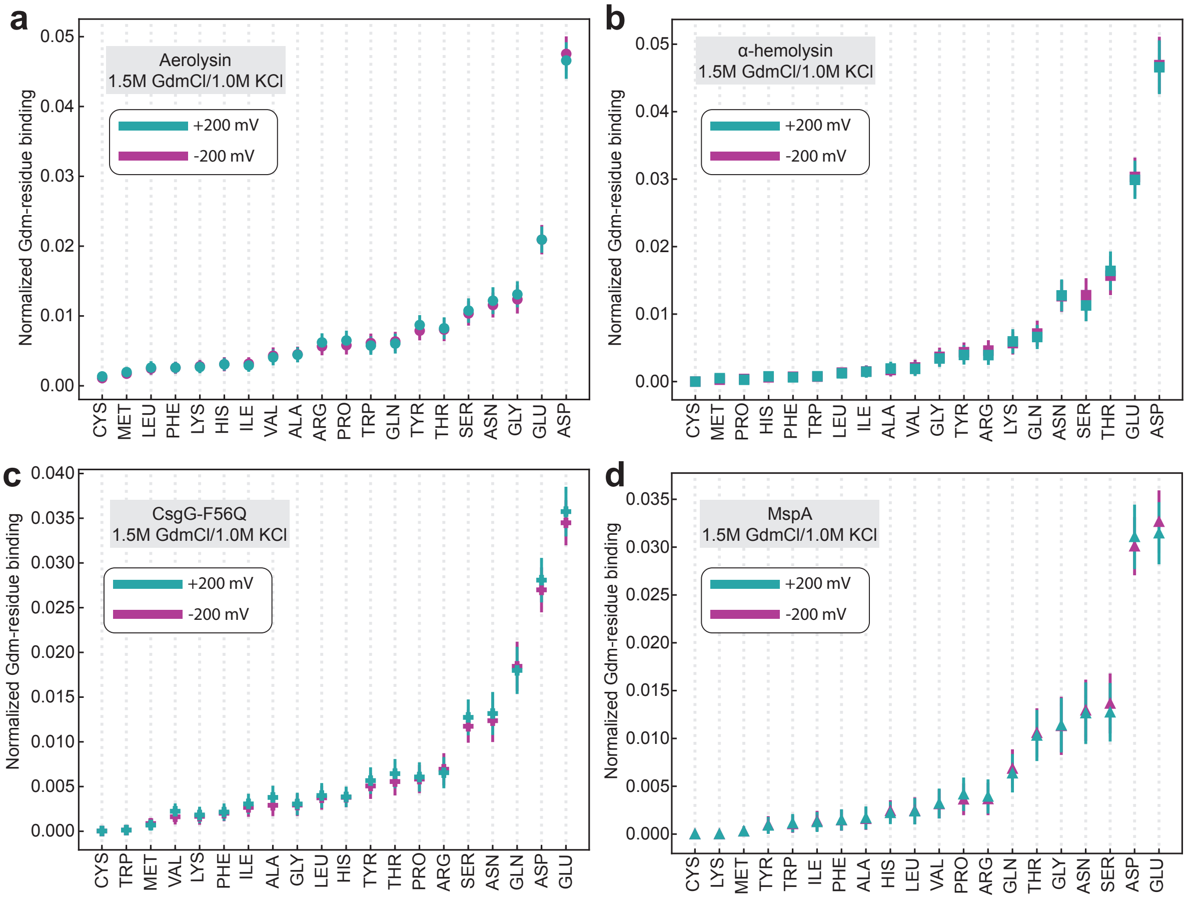

**SI Figure 9. Amino acid-specific interaction of Gdm$^+$ with biological nanopores. a–d,** Number of the residues of the specified type that were find to bind Gdm$^+$ normalized to the total number of such solvent accessible residues in MD simulations of aerolysin (panel a), $\alpha$-hemolysin (panel b), CsgG-F56Q (panel c) and MspA (panel d). Error bars show the standard error computed using 30 ns fragments of the MD trajectories. In each panel, the data are arranged in ascending order. The electrolyte and applied bias conditions are specified in each panel.

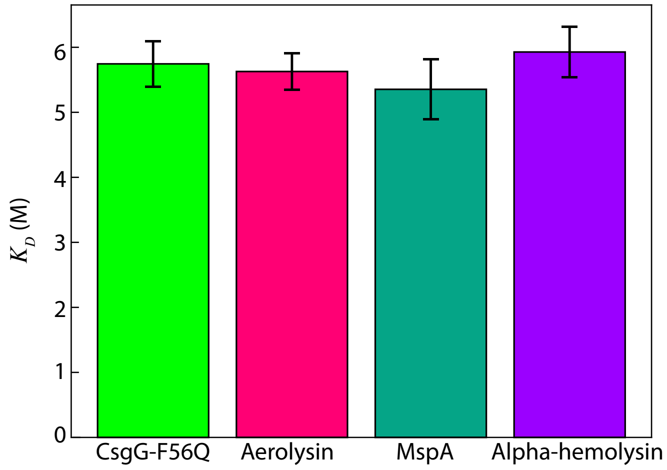

**SI Figure 10. Overall binding affinity of $Gdm^+$ to biological nanopores.** The dissociation constant ($K_D$) between $Gdm^+$ and each of the four pores was computed considering all solvent-exposed amino acids as identical binding sites. Error bars show the standard deviation of the data computed using instantaneous frames of the corresponding MD trajectory. The electrolyte and applied bias conditions are 1.5 M GdmCl / 1.0 M KCl at +200 mV.

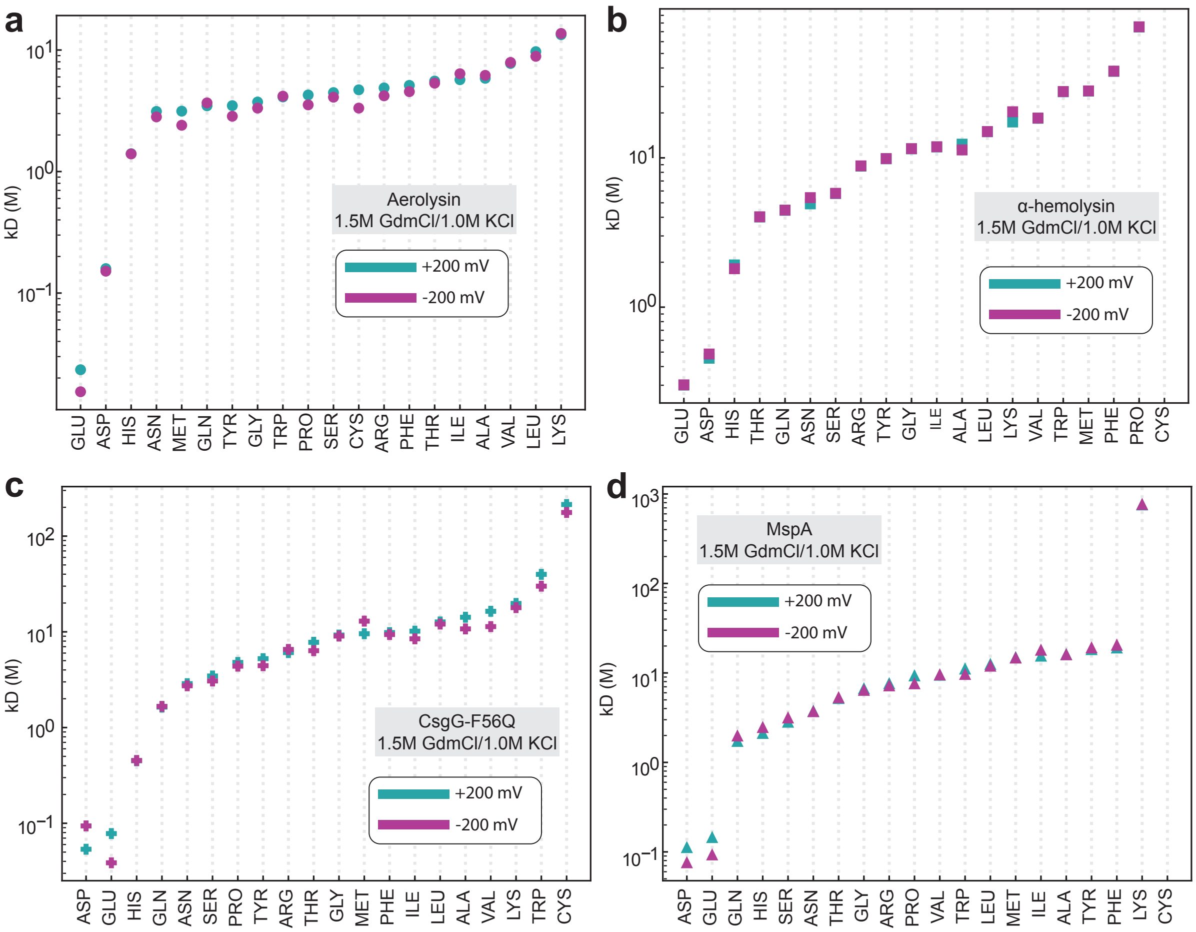

**SI Figure 11. Binding affinity of $Gdm^+$ to amino acids of the biological nanopores. a–d,** Dissociation coefficient ($K_D$) between $Gdm^+$ and each type of amino acids at the surface of aerolysin (panel a), $\alpha$-hemolysin (panel b), CsgG-F56Q (panel c) and MspA (panel d). Please note the use of a logarithmic scale for the $K_D$ values. Within each panel, amino acid types are arranged in ascending order according to the +200 mV data. The electrolyte and applied bias conditions are specified in each panel.
